# Enhancer control of promoter activity and variability *via* frequency modulation of clustered transcriptional bursts

**DOI:** 10.1101/2025.03.26.645410

**Authors:** Jana Tünnermann, Gregory Roth, Julie Cramard, Luca Giorgetti

## Abstract

Gene expression in mammalian cells is controlled by enhancers that are often dispersed across large *cis*-regulatory landscapes around a promoter. Yet how enhancers determine transcription of their target genes, and how this depends on their relative position inside a *cis*-regulatory landscape remains unclear. Here we use live-cell imaging to track the activity of a promoter under the control of the same enhancer, but inserted at different positions across a simplified regulatory landscape with minimal complexity. Combined with mathematical modeling, this reveals that RNA production from the promoter occurs in clusters of transcriptional bursts, with enhancer position controlling the frequency at which such clusters appear. This results in bursts being more frequent and occurring more uniformly across cells when the enhancer is genomically close to the promoter than when it is located at large genomic distance. Mathematical modeling further indicates that the enhancer modulates the promoter’s ability to transition from its basal transcriptional state to a regime where clusters of bursts become more frequent. Our results challenge existing models of mammalian promoter operation, and reveal that enhancer position within a *cis*-regulatory landscape critically controls the timing and variability of transcriptional output in single cells.

## Introduction

In metazoans, control of gene expression critically relies on enhancers that determine the tissue specificity and developmental timing of the transcription of large numbers of genes^1^. Enhancers can be positioned at a wide range of genomic distances from the genes that they control, and are often dispersed into *cis-*regulatory landscapes spanning several hundred kilobases around their target promoters. Despite their central role in gene regulation, how enhancers control their target genes remain elusive. Studies based on population-averaged chromosome conformation capture and RNA quantification methods have suggested that enhancers communicate regulatory information to their promoters *via* direct physical proximity enabled by the three-dimensional (3D) folding of the chromatin fiber^2–5^. However, the spatio-temporal relationship between such encounters and RNA synthesis in single cells remains unclear, especially in mammalian cells^6^.

Inside single cells, regulatory events at enhancers and promoters are inherently stochastic and dynamic due to the stochastic binding and unbinding of small numbers of transcription factors and cofactors resulting in discontinuous loading and licensing of RNA polymerase II (Pol II) complexes^7^. This generates intermittent bursts of promoter activity during which Pol II molecules are released into productive elongation followed by periods of inactivity during which no transcription is observed^8^. The physical distances between enhancers and promoters are additionally thought to fluctuate in time^9–11^, thus likely allowing molecular interactions at the enhancer-promoter interface to occur only within limited windows of time. Yet how promoter burst kinetics is regulated by stochastic interactions with an enhancer, and how this depends on the genomic location of the enhancer within the promoter’s *cis*-regulatory landscape, remains poorly understood^12^. Understanding how enhancers control burst dynamics is however fundamental to better understand how transcription levels as well as their cell-to-cell and temporal variability are encoded by genomic sequence.

We and others recently showed that at the population level, transcription levels at a promoter increase when decreasing its genomic distance from an enhancer^11,13–18^. This can be explained in terms of an increase in promoter burst frequency with increased enhancer-promoter contact probabilities, which themselves increase with decreasing genomic distance^13^, in agreement with other studies that suggested that enhancers mainly control burst frequency^19–21^. This implies that in addition to enhancer and promoter sequences themselves, their location within a *cis*-regulatory landscape and notably their reciprocal genomic distance not only control average transcription levels – but also impact the heterogeneity of mRNA production in single cells and thus the precision at which regulatory inputs can be translated into transcriptional outputs. However, direct evidence that the large-scale organization of genomic locus impacts burst dynamics in single cells is missing.

Recent studies have also questioned the validity of interpreting mammalian promoter burst kinetics in terms of simplistic two-state (ON/OFF) models of gene expression. A growing body of research supports instead the notion that mammalian promoters function as three-state models, where the promoter transitions between a repressive state (‘deep OFF’) and a permissive state (OFF), which only can transition to an active state (ON state) from which RNA is produced^20,22,23^. Some studies supported the hypothesis that mammalian enhancers play a role in modulating transitions between OFF states or between the permissive OFF and the ON state^20,22^. Yet, whether enhancers and promoters generally satisfy such 3-state kinetics, and what molecular mechanisms could account for this behavior remain unknown^24,25^.

Here we use a reductionist approach in mouse embryonic stem cells (mESC) to study the transcriptional dynamics of the Sox2 promoter while varying its genomic distance with its cognate SCR (Sox2 control region) enhancer in a controlled genomic region that minimizes regulatory and structural confounding effects. This allows us to isolate the effect of enhancer-promoter distance – and hence contact probability, which inversely scales with genomic distance – while keeping all other regulatory variables (sequence identity, biochemical composition of the enhancer-promoter interface) constant.

Quantitative analysis of live-cell microscopy experiments reveals that the genomic distance between the enhancer and the promoter modulates the promoter’s burst frequency without affecting burst duration and size. Irrespective of its genomic distance from the enhancer, burst dynamics at the promoter cannot be explained by previously proposed two-or three-state models of promoter operation, since we observe that transcription occurs in clusters of short-lived bursts separated by longer inactive periods. Interpretation of the data using mathematical modeling suggests that the promoter stochastically switches between two transcriptional regimes, one with lower and one with higher burst frequencies. Changing the genomic distance from the enhancer modulates the rate at which the promoter switches from the low-to the high-frequency regime, suggesting that interactions with an enhancer push the promoter into a state that is able to produce RNA more frequently. Finally, our data also reveal that larger enhancer-promoter distances also lead to higher cell-to-cell variabilities in the timing and number of transcription bursts, suggesting that the large-scale organization of a regulatory locus plays an important role in controlling transcriptional precision in space and time.

## Results

To study the transcriptional dynamics of a promoter under the action of a cognate enhancer with minimal confounding factors, we built on an enhancer mobilization assay that we recently established in mouse embryonic stem cels (mESC). This relies on a piggyBac transposon cassette to mobilize an enhancer and reinsert it into many random loci around an initial position within an ectopic ‘neutral’ genomic region^13^. In this assay, the mouse *Sox2* promoter drives expression of a split EGFP transcript containing a piggyBac transposon cassette harboring the Sox2 control region (SCR) enhancer^26^. This construct is integrated in a ‘neutral’ topologically associated domain (TAD) on chromosome 15 that is devoid of other active enhancers and promoters, active or repressive chromatin marks, or CTCF loops. To allow the visualization of RNA produced by this construct, we now integrated 24 MS2 stem-loop repeats^27,28^ in the 3’ untranslated region (UTR) of the split EGFP transgene (**Fig. 1a)**. When transcribed, these loops can be visualized upon binding of the bacteriophage MS2 coat protein (MCP) fused to HaloTag, which was stably expressed following random integration of the corresponding expression construct. Upon expression of the PBase transposase, the enhancer transposon cassette is excised and randomly integrated into the genome (**Fig. 1b**). Reconstitution of a functional *EGFP* transcript followed by sorting of EGFP-positive cells enables the generation of clonal mESC lines, in each of which the SCR is located in a different position within the ‘neutral’ TAD. This approach ensures all clonal mESC lines to have the same MCP-HaloTag expression levels and the enhancer-promoter genomic distance to be the only varying factor, allowing subsequent live-cell imaging data, detection of transcription start site, and assessment of burst parameters such as burst frequency, duration and size to be directly comparable between clones (**Fig. 1c**)

**Figure 1.**
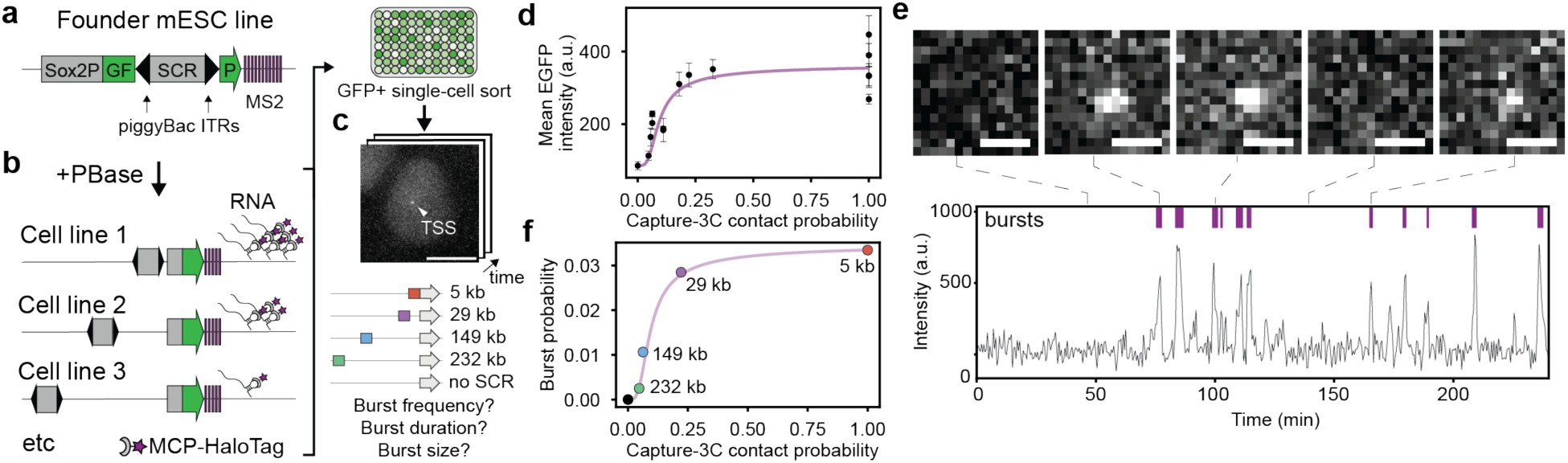
Enhancer-promoter genomic distance dictates the probability of observing transcription sites. **a.** Schematic of piggyBac-based enhancer mobilization: the Sox2 promoter (P) drives transcription of an *EGFP* gene split by a piggyBac transposon containing the SCR enhancer (E). The 3’UTR of the *EGFP* transcript carries 24 MS2 stem-loop repeats. ITR: inverted terminal repeats. **b.** Upon expression of PBase, the enhancer is excised and reintegrated elsewhere in the genome, reconstituting the *EGFP* transgene and enabling EGFP+ cells to be sorted into clonal cell lines. **c.** Live-cell imaging is used to detect transcription sites (TS) and measure burst parameters as a function of enhancer-promoter genomic distance. Scale bar: 5 µm **d.** Mean EGFP intensities assessed through flow cytometry in individual EGFP+ cell lines as a function of the contact probability between enhancer and promoter locations measured by capture-3C in Ref. ^13^. Purple line: EGFP level trendline from Ref.^13^. **e.** MS2 intensity trace from one cell with example images of TS signals (scale bar: 1 µm). Bursts were defined based on a spot being detected by the spot detection algorithm. **f.** Time-averaged probability of observing a burst at a given time point as a function of contact probability measured in capture-3C. Purple line as in panel d.

Using this modified enhancer mobilization assay, we generated 14 clonal mESC lines with ranging genomic distances of 5 to over 300 kb (**Extended Data Fig. 1a**), plus one cell line in which the SCR enhancer was reintegrated on a different chromosome from which it exerts no effect on the ectopic *Sox2* promoter **(Extended Data Table 1**). In these lines, population-averaged EGFP protein levels assessed in flow cytometry remained stable over passages (**Extended Data Fig. 1b**). EGFP levels decreased with increasing enhancer-promoter genomic distance (**Extended Data Fig. 1a)**, and scaled nonlinearly as a function of the contact probability between the promoter and enhancer insertion locations inferred from previous capture-C data^13^ (**Fig. 1d**). This trend could be well described by the nonlinear relationship that we previously measured between EGFP expression levels and contact probabilities with the same assay in the absence of MS2 stem loops (**Fig. 1d**), indicating that the stem loops do not perturb the transcriptional behavior of the Sox2 promoter under the control of the SCR.

We then selected a subset of five clonal mESC lines where the enhancer was reintegrated at genomic distances of 5, 30, 149, 232 kb (**Fig. 1c**) and on a different chromosome for further analysis. In each of them, we acquired one EGFP 10-µm z-stack followed by movies of 10-µm z-stacks every 30 seconds for 5 hours using oblique illumination microscopy (**Supplementary Video 1,2**). Average EGFP protein levels as assessed by microscopy were highly correlated to those assessed by flow cytometry, illustrating that imaged cells were representative of the population (**Extended Data Fig. 1c-d**). We then quantified the kinetics of transcription by detecting transcription sites as diffraction-limited spots in time-resolved image series. Transcriptional bursts were then defined as consecutive time frames where a transcription start site was detected (see **Methods**). Imaging 3-7 technical replicates per condition led to between 1000 and 2500 MS2 intensity tracks per condition of varying duration (example trace in **Fig. 1e, Extended Table 2**). Interestingly, the time-averaged probability of seeing a burst at a given time point revealed the same nonlinear relationship as previously reported for transcriptional levels measured by mature mRNA or protein^13^ (**Fig. 1f**), suggesting that the SCR might predominantly control the probability of initiating a burst from the *Sox2* promoter.

## Enhancer-promoter genomic distance affects inter-burst duration

We next set out to understand how quantitative burst parameters change as a function of enhancer-promoter genomic distance, and hence contact probability. For this, we calculated the survival probabilities of burst size, burst amplitude, burst duration and the inter-burst duration in the five mESC lines with different SCR positions (**Fig. 2**). To avoid confounding effects from fluorescent levels declining over the first hour of imaging (**Extended Data Fig. S2a**), we removed the first 120 frames from this analysis.

**Figure 2.**
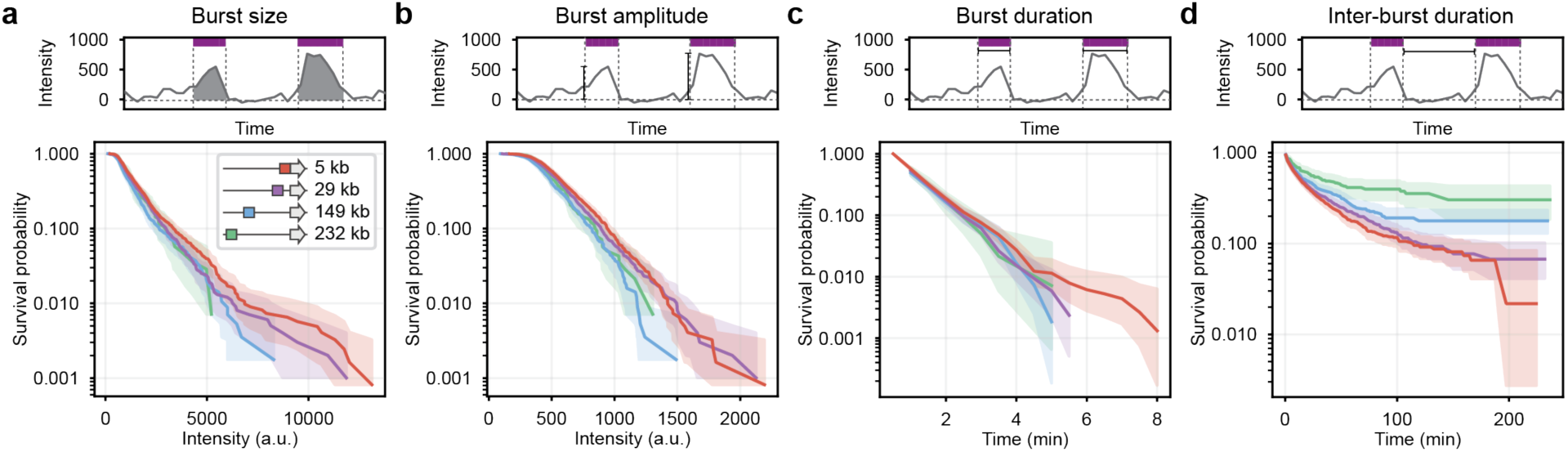
Quantification of burst parameters from live-cell imaging. Upper panels: schematics of quantified parameters. **a.** Survival probability of burst sizes. 95% confidence intervals were estimated by bootstrapping (n=10.000). **b.** Survival probability of burst amplitudes. 95% confidence intervals were estimated by bootstrapping (n=10.000) **c.** Survival probability of burst durations as calculated with the Kaplan-Meier estimator. **d.** Survival probability of inter-burst durations (Kaplan-Meier estimator).

Burst size and burst amplitude (determined as the integrated intensity and the maximum intensity, respectively, of the MS2 signal over an active transcription period) showed no notable changes across different enhancer-promoter genomic distances, as confirmed by their survival probabilities falling within the 95% confidence interval calculated from bootstrapping (10.000 repeats) (**Fig. 2a-b**) (see Methods). We only noted a small trend for burst amplitudes to be smaller for larger enhancer-promoter genomic distances (**Fig. 2a-b** and **Extended Data Fig. 2b-c**).

Given that we imaged the cells over finite windows of time, it is possible that some active or inactive periods initiated or finished outside the imaging window (‘censored’ events). To account for this effect, we estimated burst duration and inter-burst interval distributions using the Kaplan-Meier estimator (see Methods). Censoring-corrected burst durations were independent of enhancer-promoter genomic distance (**Fig. 2c, Extended Data Fig. 2d**), with individual bursts being relatively short, exhibiting a median survival time of approximately one and a half minutes. However, inter-burst durations substantially increased with increasing enhancer-promoter genomic distance (**Fig. 2d, Extended Data Fig. 2e**), with median survival times ranging from 15 minutes for the two closest enhancer-promoter genomic distances (5 kb and 30 kb, respectively) up to 30 minutes for the longest enhancer-promoter genomic distance (232 kb), each displaying wide distributions from one minute to hundreds of minutes. Interestingly, the probability that inter-burst events lasted more than four hours increased with increasing enhancer-promoter distance, ranging from 2% at 5 kb to 30% at 232 kb (**Fig. 2d**). This reflects the observation that the fraction of cells that never burst in four hours also increased with increasing distance (**Extended Data Fig. 2f**).

To ensure that these measures were not artifacts of our image analysis pipeline, we manually curated 10% of the live-cell imaging data from the two clones with enhancer-promoter genomic distances of 5 kb and 149 kb. Comparing the survival probabilities from the manually annotated data with the automatic image analysis pipeline revealed no differences in distribution or quantities, thus validating our data processing method (**Extended Data Fig. 2g-n**).

Together, these data indicate that the genomic distance with its enhancer modulates the burst frequency of the Sox2 promoter, with no effect on burst duration and only very minor effects on burst amplitude and size.

## Burst frequency and its variability across cells scale nonlinearly with contact probability

In line with these findings, we observed that the Sox2 promoter’s burst frequency, defined as the average number of active transcription periods per hour, fully recapitulated the nonlinear relationship between transcriptional output and contact probabilities that we previously observed^13^ (**Fig. 3a**; see **Fig. 1d, f**). As we previously noted however (see **Fig. 2d**), inter-burst durations are widely distributed in each clone. To illustrate this point we considered two clones with enhancer-promoter genomic distances of 5 kb or 149 kb whose average burst frequencies were approximately 7 and 1 burst over the course of four hours, respectively. Binarized transcription traces from individual cells revealed substantial cell-to-cell variability in the number of bursts, with many cells bursting way more than seven times with the enhancer at 5 kb distance, whereas many cells showed no bursts at all with the 149 kb away (**Fig. 3b,c**). Quantification of this effect using the coefficient of variation of burst frequency showed that genomic proximity of the enhancer to the promoter correlates with reduced cell-to-cell variability in the number of bursts produced in a defined amount of time (**Fig. 3d**). This indicates that enhancer-promoter genomic distance controls the reliability of promoter output in time.

**Figure 3.**
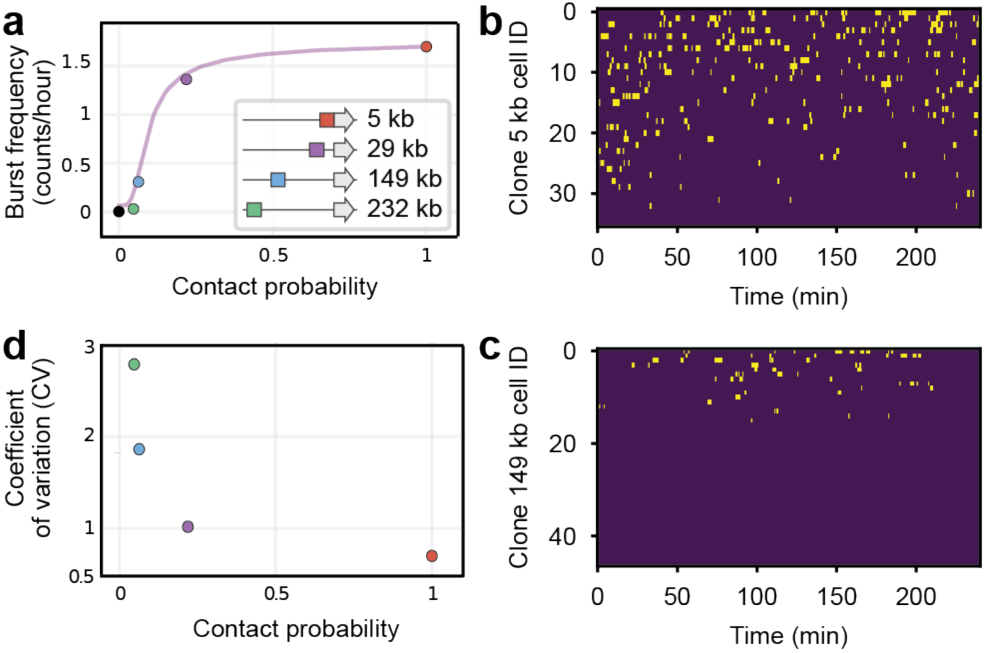
Enhancer-promoter distance modulates burst frequency and its variability across cells. **a.** Burst frequency (number of bursts per hour) as function of enhancer-promoter contact probability (mean tracks lasting four hours). Purple trendline from Ref.^13^. **b-c.** Binarized intensity traces lasting at least 4 hours for clonal cell lines with enhancer-promoter genomic distances of 5 kb **(b)** and 149 kb **(c)**. Every row corresponds to one cell; traces are sorted by the number of bursts in the trace. **d.** Cell-to-cell variability of burst frequency as function of contact probability quantified via the coefficient of variation of burst frequencies.

## Transcriptional bursts occur in clusters

Contrary to the predictions of a simple two-state model of promoter operation, survival probabilities of inter-burst durations (**Fig. 2d**) are not single exponentials. Quantitative analysis of survival probabilities revealed instead that they can be best described by a weighted sum of 3 exponential distributions (**Fig. 4a** and **Extended Data Fig. 3a**). The lifetimes of the exponentials vary between clones within 3-7, 24-52, and >164 minutes (**Extended Data Fig. 3c**). Fitting with one or two exponentials failed to reproduce the full behavior of the curve (**Fig. 4b** and **Extended Data Fig. 3a-d**). These multiple timescales over which no burst is observed suggest the existence not of a single, but rather multiple OFF states with distinct lifetimes.

**Figure 4.**
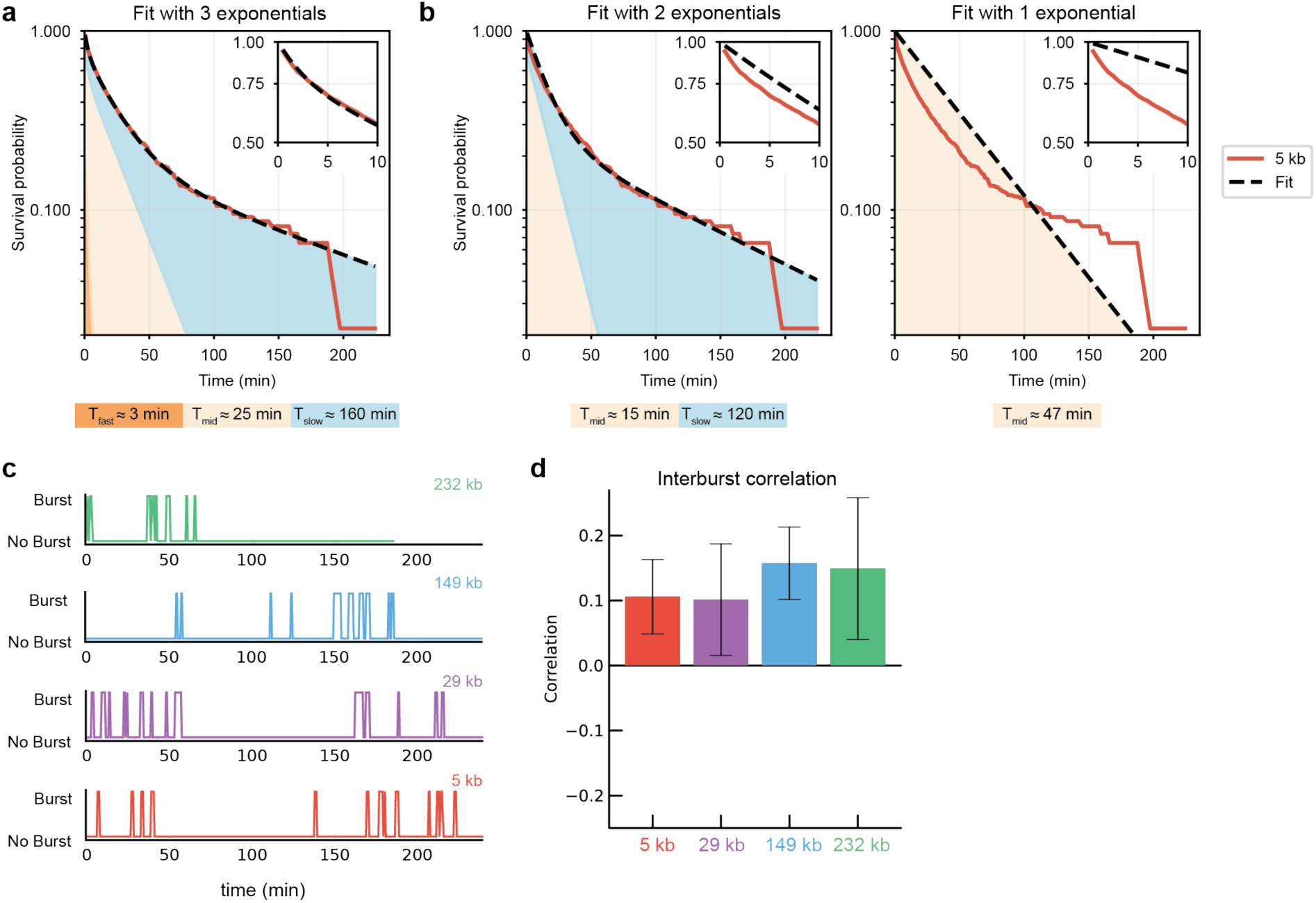
Inter-burst durations are distributed over three timescales. **a.** Best fit of the three-exponential model to experimental survival probabilities of the inter-burst durations in the cell line where the SCR is located 5 kb away from the promoter. The best fit curve (black dotted line) is the weighted sum of three exponential curves represented in colored areas from the highest rate (orange) to the lowest rate (blue). The timescale of each exponential curve (1/rate) is shown in the bottom. Insets: zoom of the plots over the 0-10 minutes range. **b.** Same as panel a) but for the two-exponential model (left) and single exponential model (right). **c.** Examples of binarized traces illustrating clusters of bursts. Color corresponds to cell line identity as in Fig. 2. **d.** Pearson correlation between the durations of consecutive inter-burst events, measured in the cell lines shown in Fig. 1c. Mean ± s.d based on 2000 bootstrap runs over the set of tracks.

Interestingly, visual inspection of the tracks suggested that bursts often occur in groups of consecutive events separated by larger periods of inactivity (**Fig. 4c**). Quantitative analysis of the tracks revealed indeed that the survival probability of inter-burst periods had inflated frequencies of both short and long inter-burst intervals compared to a single exponential curve (**Extended Data Fig. 3d**). This is a signature of clustered patterns in a stochastic sequence of events^29^, indicating that bursts from the promoter do not occur randomly, but rather in clusters.

We also found that the correlation between consecutive inter-burst periods was small but positive, indicating that short inter-burst times tend to be followed by short inter-burst times and vice versa for long inter-burst times (**Fig. 4d**). Positive correlation between consecutive inter-burst periods suggests that the promoter operates in at least two transcriptional ‘regimes’: a ‘high-frequency’ regime in which the promoter is transcribed in clusters of bursts, that is short inter-burst periods are followed by other short inter-burst periods; and a ‘low-frequency’ regime in which the promoter only bursts sporadically (long inter-burst periods followed by long inter-burst periods) (**Extended Data Fig. 4a**). If this wasn’t the case, i.e. if the promoter operated in a single frequency regime, or two regimes of which one never bursts – then correlations would be zero (**Extended Data Fig. 4b-c**). Together, these observations indicate that transcriptional dynamics in this system cannot be described by models of promoter operation that consider a single frequency regime, and notably by previously proposed two-or three-state models with a single ON state and one or multiple OFF states.

## Modeling the multi-state kinetics of promoter operation

We next wondered which stochastic model of promoter operation could account for the ensemble of our experimental observations. The simplest double-regime model that predicts positive correlations, as well as three distinct timescales in the inter-burst durations as observed in our data is a stochastic transition between two separate 2-state regimes. In this model, each of the two regimes has an ON state from which RNA synthesis can be initiated, and an OFF state that is not transcribed; the two regimes only differ in their OFF-to-ON transition rates, which is small in the ‘low-frequency’ regime and larger in the ‘high-frequency’ regime (**Fig. 5a**). In such a model, the longest and intermediate inter-burst timescales arise from OFF->ON transitions in the low-frequency and high-frequency regimes, respectively. The fastest timescale only arises if the rate at which transcription is initiated in the ON state(s) is small compared to the rate at which RNA is released once it has transcribed, thus creating a gap in the fluorescence signal (**Fig. 5b**).

**Figure 5.**
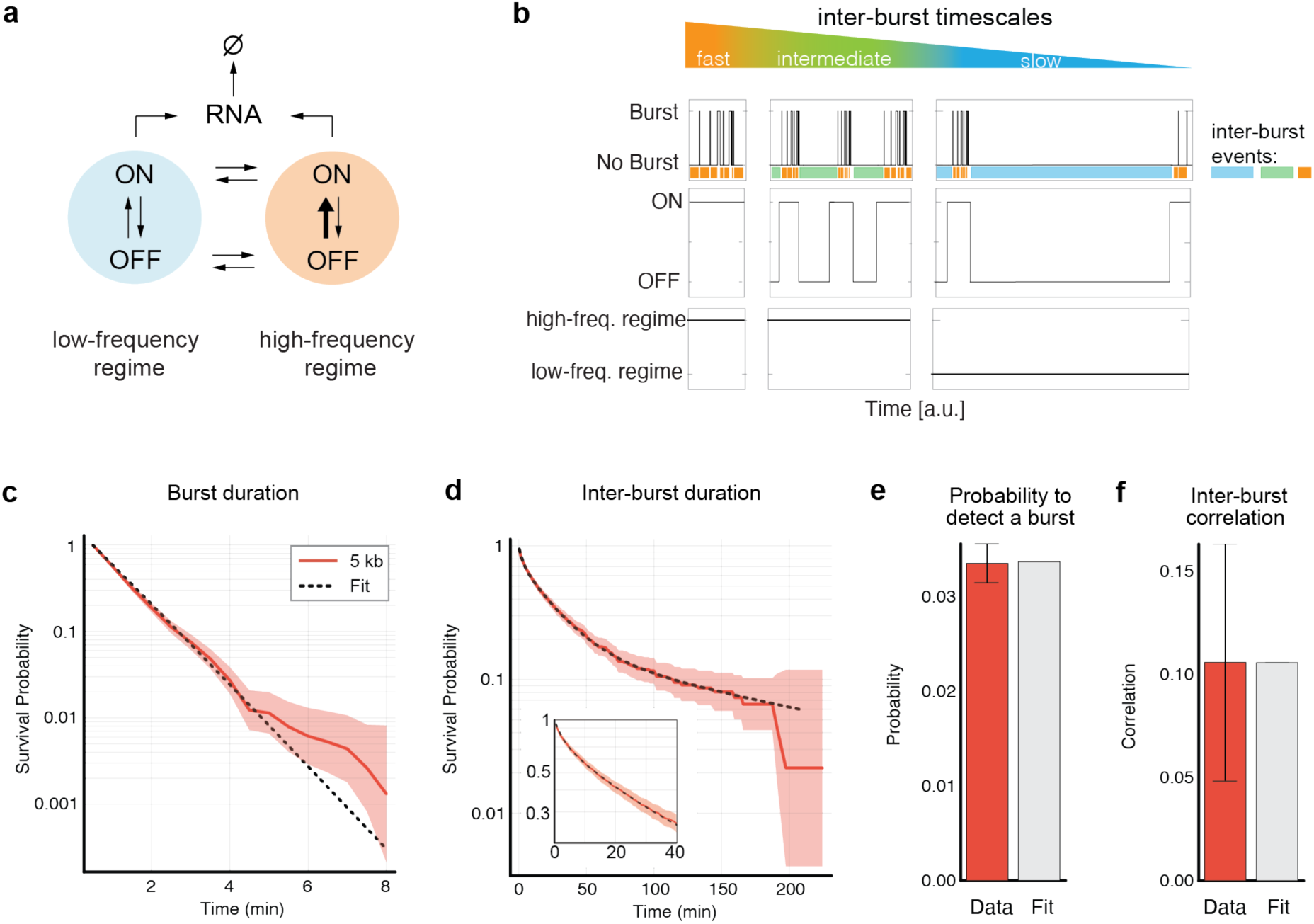
A double-regime two-state model of transcriptional dynamics. **a.** Diagram of the model. The promoter operates in a ‘low-frequency’ regime with a small ON rate and can transiently enter a ‘high-frequency’ regime with a higher ON rate (bold arrow). **b.** Schematics of the dynamics of transcription bursts (ongoing transcription), promoter ON/OFF states, and promoter low/high regimes, illustrating the three timescales of the inter-burst durations. **c-f.** Best fit of the double-regime two-state model to the observed survival probabilities of burst durations **(c)**, inter-burst durations **(d)**, steady-state probabilities to detect a burst (**e**), and the Pearson correlation of consecutive inter-burst periods **(f)** for the cell line where the SCR was located 5 kb from the promoter. The confidence band in panes (c) and (d) is +/-the pointwise standard error of the log of the survivor function, computed using Greenwood’s formula. In (e) and (f), data are mean ± s.d, based on 2000 bootstrap runs over the set of tracks.

To test whether this model can quantitatively reproduce our experimental observations, we first fit it to the data of a single clone (enhancer-promoter distance = 5 kb). Specifically, we simultaneously fit the model to four observables: the survival probabilities of i) burst and ii) inter-burst durations, iii) the steady-state probability to observe a burst, and iv) the correlation between consecutive inter-burst periods (see **Supplementary Information**). Given the large parameter space and the computational cost of tracking all stochastic events in simulated trajectories, direct simulation-based optimization was impractical. To overcome this, we developed a theoretical framework that provides analytical expressions for these observables, allowing for efficient parameter exploration (see **Supplementary Information**). The model reproduced well these four experimental observables (**Fig. 5c-f**). Best agreement occurred with an average burst duration of 1.2 minutes, an average inter-burst duration of 4 hours in the low-frequency regime, and 21 minutes in the high-frequency regime (**Extended Data Fig. 4c**). The high-frequency regime lasts on average 4.7 hours and produces 3.6 bursts per hour. The low-frequency regime lasts on average 7 hours and produces 0.3 bursts per hour. In steady-state, 40% of the cells are in the high-frequency regime, resulting in an overall average inter-burst duration of 49 minutes and an overall average of 1.7 bursts per hour (**Extended Data Fig. 4d**).

We then asked if the data from the other clones could be explained under the stringent hypothesis that enhancer-promoter genomic distance modulates only one rate in the model. In principle, all rates could affect inter-burst duration while keeping similar burst duration as observed in the experiments (**Fig. 2c-d**). Thus for all rates, we simultaneously fit the model to the survival probabilities of the inter-burst duration of all other clones (enhancer-promoter distance = 29 kb, 149 kb, 232 kb), when only the rate of interest can differ between clones and all other parameters are fixed to the best-values of the 5kb clone; we then asked if the best-fit parameters could predict the observed inter-burst correlations (Model analysis in **Supplementary Information**). Three scenarios (changing ON rate in the low-frequency regime, changing OFF rate, and changing initiation rate) were not able to reproduce the inter-burst durations in the first place (**Extended Data Fig. S5a-c**). Two other scenarios (changing ON rate in the high-frequency regime, and changing high-to-low regime switch rate) reproduced inter-burst durations well, but failed at predicting both the trend and amount of inter-burst correlations (**Extended Data Fig. 5d-e**). Interestingly, the only scenario where the low-to-high regime switch rate depends on enhancer-promoter distance was able to fit the inter-burst durations and simultaneously predict both the trend and amounts of correlation between consecutive inter-burst times (**Fig. 6b-c**). Best agreement occurred with an average inter-burst duration of 1.0 (enhancer-promoter distance = 29 kb), 1.6 ( distance = 149 kb), and 3.1 hours ( distance = 232 kb), resulting in a burst frequency of 1.4, 0.9, and 0.4 bursts per hour (**Extended Data Fig. 4d-f**).

**Figure 6.**
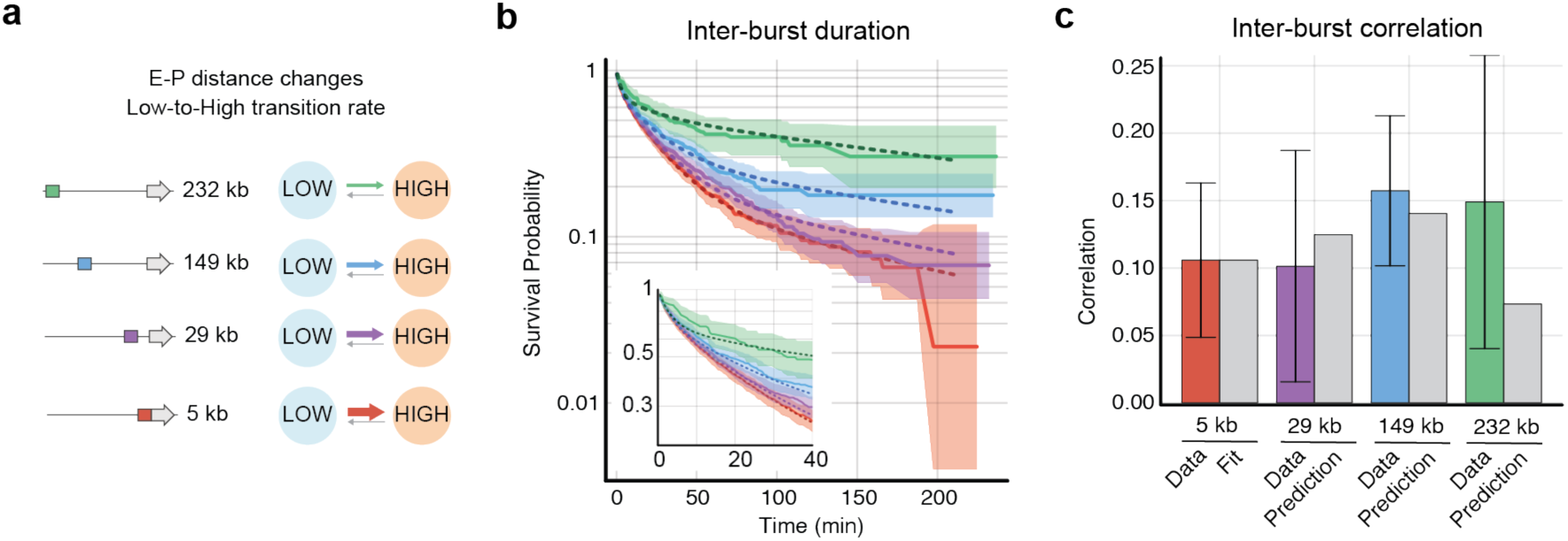
Enhancer-promoter distance controls the rate to switch from the low-frequency to the high-frequency regime. **a-b**. Survival probabilities of inter-burst duration in cell lines with the SCR at different distances from the promoter can be predicted (purple, blue, and green dotted lines) from the model fitted to the cell line where the SCR was located at 5 kb (red dotted line), with a modified low-to-high regime rate. Data is similar to Figure 2d. **c.** Best-fit and predictions of the Pearson correlation between consecutive inter-burst periods. Experimental data same as in Fig. 4d.

Taken together, our results are compatible with a model in which the promoter spends a large fraction of time in a low-frequency bursting regime, in which the rare OFF->ON transitions give rise to the longest inter-burst timescale. Occasionally, the promoter switches to a high-frequency regime, whose OFF->ON transitions give rise to the observed intermediate inter-burst timescale (∼25 min). The fastest of the three inter-burst timescales (∼3 minutes) arises instead from the initiation rate in the ON state being small compared to the rate at which the RNA is released after being transcribed (**Fig 5b**). This leads to disappearance of the fluorescence signal because no RNA synthesis was initiated in the time required to elongate and release the preceding RNA. Our analysis also indicates that a substantial number of bursts are in fact single transcription events, which occur one after another during a single ON state. When the promoter switches to the high-frequency regime, such groups of bursts occur more frequently. Interestingly, the rate at which the promoter switches to the high-frequency regime is controlled by the enhancer, which thus controls the promoter’s effective burst frequency by modulating the frequency at which it produces clusters of bursts.

## Discussion

Our study provides quantitative measurements of promoter burst kinetics when varying its genomic distance from an enhancer, over a range that is representative of those observed within mammalian genomes (from few to hundreds of kb). By only modifying the position of the enhancer within a ‘neutral’ genomic region with minimal structural and regulatory complexity, we could isolate the effect of genomic distance from other confounding factors such as enhancer-promoter biochemical compatibility and influence from additional regulatory sequences, which may confound interpretation of results when enhancer sequences are manipulated at endogenous gene loci.

In line with previous reports that enhancers may mainly affect burst frequency^11,19,20,22,30^, we observe that the genomic distance between the *Sox2* promoter and its SCR enhancer modulates the time between consecutive bursts, without any noticeable changes in burst size or duration (**Fig. 2**). In contrast to previous studies, however, we further reveal that bursts from an enhancer-driven promoter occur in clusters (**Fig. 4c** and **Extended Data Fig. 3e**). This leads to the appearance of correlated inter-burst durations, which are distributed over three well-separated timescales (**Fig. 2, 4**).

This kinetic behavior is not compatible with any of the existing models of promoter operation which consider a single burst regime, and specifically two-or three-state models of promoter operation^20,22,23^. Instead, we show that it can be generated by an alternative simple model in which the promoter can switch between a low-frequency and a high-frequency two-state regimes. Testing this model against experimental perturbation of enhancer-promoter distance suggests that the enhancer controls the frequency at which consecutive clusters of bursts appear (**Fig. 5**), rather than the frequency at which single bursts occur within a cluster. We note that one key feature of such model is that transcription is initiated inefficiently from the ON state, in line with recent interpretations of transcriptional kinetics in *Drosophila*^24^.

The structure of the model, which we inferred purely from the analysis of transcriptional dynamics in single cells, strongly resembles – and thus supports – the structure of a model we previously proposed to describe population-based measurements in the same experimental system^13^. In Ref.^13^ we made the explicit hypothesis that the switch between the low-frequency and the high-frequency regimes is triggered by stochastic enhancer-promoter interactions *via* one or more reversible regulatory steps, which allow the promoter to enter the high-frequency regime once they are completed. Although the model we describe here does not explicitly consider enhancer-promoter interactions and their effect on the promoter, the finding that the low-to-high-regime transition rate depends on enhancer-promoter genomic distance (and thus on their contact probability) still suggests that enhancer-promoter interactions determine regime switching (**Fig. 6**).

We note that stochastic transitions from the low-to the high-frequency regime were inferred to occur on average only once every 7, 10, 25 and 151 hours when the enhancer is located at 5, 29, 149 and 232 kb, respectively (cf. low-to-high rates in **Extended Data S4d**). This suggests that interactions with the enhancer might only rarely translate into a detectable regulatory effect at the promoter, especially when the enhancer is located tens to hundreds of kb away. This is in line with our observation that substantial numbers of cells do not burst over the course of 4 hours (**Extended Data S2f**) and with other recent measurements on endogenous genes in mESC^20^. The experiments we report in this study do not give access to the dynamics of enhancer-promoter distances. It is nevertheless possible to compare the burst frequency and regime transition rate we measured at 149 kb distance to the frequency of events of physical proximity that we previously measured between Tet operator (TetO) and Lac operator (LacO) arrays located 150 kb apart in the same ‘empty’ TAD^9^. On average, these sequences returned into physical proximity (defined as a range of ∼150 nm) 3.5 times an hour, which is around three times more frequent than the average burst frequency we measured here (∼one burst every three hours, **Fig. 3a**), and 100 times more frequent than inferred promoter transitions from the low-to the high-frequency regime (**Fig. 3a**). If the dynamics of enhancers-promoter looping resemble those of non-regulatory DNA (i.e. LacO and TetO), these figures would suggest that the majority of such proximity events might either represent non-functional interactions, or consecutive proximity events leading to one productive regulatory event *via* transitions through intermediate rate-limiting regulatory steps^13,31^. Alternatively, only a subset of proximity events (e.g. those that are longer than a certain time^32^) might allow the transition from the low-to the high-frequency promoter regime, which is then maintained via molecular processes that do not depend on the permanence of a contact.

In addition to controlling population– and time-averaged transcription levels, this also results in different levels of heterogeneity across cells in the timing and number of bursts produced in a fixed time window – with enhancers located close to the promoter driving larger, but also more similar numbers of bursts per unit time compared to enhancers located at larger genomic distances. This suggests that the large-scale organization of a *cis-*regulatory landscape may have a substantial effect on the accuracy and robustness^33^ of transcriptional programs. It could thus be expected that genes involved in fast and reproducible transcriptional responses might benefit from genomically proximal enhancers, whereas longer-range enhancers might be tolerated (or favored) when higher levels of heterogeneity in the timing or absolute level of expression are allowed. Interestingly, a similar pattern was recently observed in neurons^34,35^.

Finally, we note that bursts from the Sox2 promoter in this experimental system have a similar duration, they are rarer than those that were previously detected at the endogenous Sox2 locus^36,37^, even when the enhancer is in closer genomic proximity than the endogenous SCR. This might be due to the absence of the CTCF loops that connect the endogenous Sox2 promoter and SCR^38^, the lack of additional enhancers^17,39^, or additional sequence features that facilitate promoter operation at the endogenous gene^40^.

Taken together, our data uncover quantitative principles by which a distal enhancer controls transcription and its variability in single cells, and provide a quantitative framework for future studies to address mechanistically the interplay between enhancer-promoter interactions and transcription bursting.

## Acknowledgments

We thank the Giorgetti lab, Tineke Lenstra and Wim Pomp for thoughtful discussions on experiments and data analysis and feedback on the manuscript; Jeff Chao and Franka Voigt for reagents and advice on MS2 stem-loops; Tim-Oliver Bucholz, Sabine Reitner and Laurent Gelman from the FMI Facility for Advanced Image Analysis and Microscopy (FAIM) for discussions on image analysis and technical support; and Ralph Grand and Dirk Schübeler for kindly donating plasmid templates. J.T. was supported by a Marie Skłodowska-Curie Innovative Training Network (grant no. 813282 ‘PEP-NET’). Research in the Giorgetti laboratory is funded by the Novartis Foundation, the European Research Council (grant no. 759366, ‘BioMeTre’) under the European Union’s Horizon 2020 research and innovation program, and the Swiss National Science Foundation (grants no. 310030_192642 and TMCG-3_213782).

## Author contributions

J.T. conceived the study with L.G. and performed all experiments with help in cell culture from J.C., and conducted image and data analysis. G.R. performed modeling and data analysis. J.T., G.R. and L.G. wrote the paper.

## Data availability

Image analysis code used in this study is available at https://github.com/fmi-basel/ggiorget_ms2analysis

## Methods

### Culture of embryonic stem cells

All cell lines are based on the E14Tg2a parental mESC line (karyotype 19, XY, 129/Ola isogenic background). Cells were cultured on gelatin-coated culture plates in Glasgow Minimum Essential Medium (Sigma-Aldrich, G5154) supplemented with 15% fetal calf serum (Eurobio Abcys), 1% L-Glutamine (Thermo Fisher Scientific, 25030024), 1% Sodium Pyruvate (Thermo Fisher Scientific, 11360039), 1% MEM Non-Essential Amino Acids (Thermo Fisher Scientific, 11140035), 100 μM β-mercaptoethanol (Thermo Fisher Scientific, 31350010), 20 U/ml leukemia inhibitory factor (Miltenyi Biotec, premium grade) in 8% CO2 at 37 °C. For enhancer mobilization and subsequent experiments, the culture medium was additionally supplemented with 2i inhibitors (1 μM MEK inhibitor PDO35901 (Axon, 1408) and 3 μM GSK3 inhibitor CHIR 99021 (Axon, 1386). Cells were tested regularly for mycoplasma contamination and no contamination was detected.

### Targeting vectors for the integration of 24x MS2 stem-loops

The founder cell line carrying the piggyBac transgene and MS2 repeats is based on the founder cell line used in Figure 1 of ref. This cell line enables the piggyBac-mediated mobilization of the full-length Sox2 control region (SCR) enhancer (lacking its internal CTCF site^13^), inside a ‘neutral’ TAD on chromosome 15 from which two CTCF sites were deleted. The transgene was inserted into position chr15:11717617 (mm10) in Ref.^13^.

We modified this cell line by inserting a 24x MS2 stem-loop cassette in the 3’UTR of the split *EGFP* transcript. To knock-in the MS2 repeats^28^ we used three plasmids: a targeting vector, a gRNA vector to linearize the targeting vector and a gRNA vector targeting the 3’UTR of the *EGFP* transcript at its genomic locus. The targeting vector has a pUC57 backbone (Genewiz) and contains 24x MS2 repeats flanked by homology arms and a bacterial gRNA sequences for linearization of the palsmid.To linearize the targeting vector we used a modified pC2P plasmid including the bacterial gRNA sequences and a Cas9-P2A-puromycin cassette, which was kindly donated by Dirk Schübeler^41^. The gRNA sequence for the knock-in of the MS2 cassette (targeting a position 3-prime of the splitGFP gene and in front of the SV40 polyA sequence) was designed using an online tool (https://eu.idtdna.com/site/order/designtool/index/CRISPR_SEQUENCE) and purchased from Microsynth AG. This gRNA sequence was cloned into the PX459 plasmid (Addgene, cat# 48139) using the BsaI restriction site. Primers used are listed in **Extended Data Table 3**.

### Generation of a founder cell line carrying 24x MS2 stem-loops

The founder line carrying the piggyBac transgene and MS2 repeats was generated by transfecting 1.5×10^6 cells with 1.5 μg MS2 targeting vector, 750 ng PX459 (splitGFP_gRNA/Cas9) and 300 ng PX459 (linearization_gRNA/Cas9) using Lipofectamine3000 (Thermo Fisher Scientific, L3000008) according to the manufacturer’s guidelines. 24 hours after transfection, 1 µg/ml of puromycin (InvivoGen, ant-pr-1) was added to the medium for 24 h. Cells were cultured in standard medium for an additional 4 days after which single cells were isolated in 96-well plates. Sorted cells were kept for 2 days in medium supplemented with 100 μg/µl primocin (InvivoGen, ant-pm-1) and 10 µM ROCK inhibitor (STEMCELL Technologies, Y-27632). 10 days after sorting, plates were duplicated and genomic DNA extracted on-plates by lysing cells with lysis buffer (100 mM Tris-HCl pH 8.0, 5 mM EDTA, 0.2% SDS, 50 mM NaCl, 1 mg/mL proteinase K (Macherey-Nagel, 740506) and 0.05 mg/mL RNase (Thermo Fisher Scientific, EN0531)) and subsequent isopropanol precipitation. Individual cell lines were analyzed using genotyping PCR (genotyping primers listed in Extended Data Table 3) and correctly genotyped cell lines were expanded. Correct insertion was verified by Sanger sequencing of the genotyping PCR result covering the whole insert.

Stable integration of stdMCP-stdHaloTag was achieved via lentiviral transduction (gifted by Jeffery Chao). 0.5×10^6 cells were transfected using Polybrene Infection/Transfection Reagent (Sigma-Aldrich, TR-1003) according to the manufacturer’s guidelines and 10 days later sorted for HaloTag expression into single cells as described above. After 10 days, one-third of the cells were replated onto Cellvis Glass Bottom Microplates (96-well, Cellvis, P96-1.5H-N) and clonal lines were screened by microscopy for a good signal-to-noise ratio (SNR).

### Mobilization of the piggyBac-enhancer cassette

Mobilization of the enhancer cassette was performed as previously described^13^ with minor adjustments. In short, cells were cultured in medium additionally supplemented with 2i inhibitors. PiggyBac-enhancer cassette mobilization was performed in 12 independent transfections by transfection of 3×10^5^ cells with 1µg of an expression vector for the PBase transposase (gifted by Jesse Owens^42^). EGFP+ cell lines were then isolated using FACS in 96-well plates and expanded for DNA extraction, flow cytometry and freezing.

### Mapping piggyBac-enhancer insertion sites in individual cell lines

PiggyBac integration sites were mapped using Splinkerette PCR after DNA extraction as described previously^13^. Splinkerette primers are listed in **Extended Data Table 3**. A modified script to analyze resulting Sanger sequencing reads can be found on GitHub: https://github.com/fmi-basel/ggiorget_ms2analysis.

### Flow cytometry

EGFP levels of individual cell lines were measured on-plate using the BD LSRII SORP flow cytometer equipped with BD High Throughput Sampler (HTS). For each clone, measurements were performed in triplicates and mean EGFP fluorescence intensities were calculated using FlowJo. All three replicates were averaged.

After expansion, before cell fixation and in parallel to live-cell imaging, EGFP levels of individual cell lines were measured in-tube using the BD LSRII SORP flow cytometer.

### Live-cell imaging

35-mm glass-bottom dishes (Mattek, P35G-1.5-14-C) were coated with 1–2 μg/mL laminin (Sigma-Aldrich, L2020) in PBS at 37 °C overnight. The day before imaging, 0.8-1×10^6 cells were seeded in Fluorobrite DMEM (Gibco, A1896701) supplemented with culture medium containing 2i inhibitors (see above) and incubated at 37 °C, 8% CO2.

Right before imaging, Halo-tagged MCP was labeled with Halo-Ligand-JaneliaFluor 549 (Tocris Bioscience, custom synthesis based on Cat. No. 6503). To this aim, medium was exchanged for new medium containing 100 nM Halo-Ligand for 20 min. Subsequently, cells were washed three times with PBS and fresh Fluorobrite medium was added.

Cells were imaged with a Nikon Eclipse Ti-E inverted widefield microscope equipped with a Total Internal Reflection Microscopy iLAS2 module (Roper Scientific), a Perfect Focus System (Nikon), a motorized Z-Piezo stage (ASI), Evolve 512 Delta EMCCD cameras with a pixel size of 16×16 μm^2 (Photometrics) and a CFI APO TIRF 100×/1.49 NA oil immersion objective (Nikon). The following laser lines were used: 200 mW Toptica iBEAM SMART laser for 488 nm, and 200 mW Coherent Sapphire laser for 561 nm. The microscope was operated in oblique illumination mode using the Visiview software (Visitron). An enclosed microscope environmental control set-up (The BOX and The CUBE, Life Science Instruments) was used at 37 °C and 8% CO2.

Before acquisition of movies, one z-stack (21 planes, spacing of 300 nm) of EGFP levels was recorded using 10% 488 nm laser power, an exposure time of 150 ms and a gain of 50. To image MS2 signals, series of z-stacks (21 planes, spacing of 300 nm) were acquired every 30 s using 4% 561 nm laser power, an exposure time of 100 ms and a gain of 100.

### Flatfield correction

Before the acquisition of movies, Mattek dishes filled with 20 nM Alexa Fluor™ 488 Hydrazide (ThermoFischer, A10436) or 100 nM Alexa Fluor™ 568 Hydrazide (ThermoFischer, A10437) were imaged using the same acquisition settings (see above). 200 z-planes with 300 nm spacing were acquired and subsequently 21 planes selected close to the cover slip. Three z-stacks were averaged per day. Dark images were acquired by setting laser power to 0%, 500 planes acquired and averaged. Flatfield correction was performed using following equation as step of image processing (see below):

(Image – darkimage) x mean(flatfieldimage-darkimage)/ (flatfieldimage-darkimage)

### Image processing of live-cell movies

Raw images were z-projected (mean projection for 488 nm channel (EGFP), max projection for 561 nm channel (MS2)) using Fiji and further processed using custom Python code. All code is available on GitHub at https://github.com/fmi-basel/ggiorget_ms2analysis.

Individual cells were segmented using StarDist and the ‘2D_versatile_fluo’ model^43,44^ and tracked over time using linear assignment problem (LAP) tracker with no gaps allowed, a minimal cell size and a minimal track length of 10 frames.

Active transcription spots were detected using Trackpy (thresholding after a band pass filtering (https://github.com/soft-matter/trackpy)) and subsequently filtered by size and intensity to remove spurious detections.

Spots were linked over time in individual cells and when no spot was visible between two bursts, the position of the promoter was linearly interpolated taking cell lateral movement and relative deformation into account. If two spots were detected in the same cell at the same time, the brighter spot was chosen for linking.

Spot intensity was read-out after projected images were flatfield corrected (see above) using a circular mask (7 pixel diameter). The local background intensity was read out using a ring-shaped mask (1 pixel thickness) around the spot mask separated by a 4-pixel gap. The global background intensity of the complete cell was read out by using the cell mask.

To construct intensity traces, spot and local background intensities were bleach corrected by dividing the normalized global background intensity. Subsequently, the local background intensity was subtracted from the spot intensity.

To segment transcriptional active/inactive times, information on spot detection was used. Whenever a spot is detected it is defined as transcriptional active time. To avoid confounding effects from fluorescent levels declining over the first hour of imaging (**Extended Data Fig. S2a**), the first hour of movies was excluded from analysis. To limit splitting inter-burst times by one-frame events, we removed 1-frame bursts from binarized traces.

Burst size was determined as the integrated intensity for the MS2 signal over an active transcription time. Burst amplitude was calculated as the maximum intensity of an active transcription time.

### Time-averaged probability to detect a burst in live-cell imaging

To calculate the time-averaged probability of seeing a burst, binarized MS2 intensity traces (transcriptional active/inactive times) were used. Per biological sample (cell line), the probability to see a transcriptional active time per time point was calculated and subsequently averaged over time.

### Survival probabilities calculated with Kaplan-Meier estimator

To account for transcriptional active/inactive times that start or end outside the imaging window (‘censored’ events), Kaplan-Meier estimator^45^ was used to calculate survival probabilities of burst and inter-burst durations. This non-parametric method utilizes both uncensored (start and end within the imaging window) and right-censored data (start within the window). Tracks where no burst was detected (censored on both sides) were not included in the Kaplan-Meier calculation.

From binarized MS2 intensity traces, we calculated inter-burst and burst durations for both non-censored and right-censored events. The survival function S(t), representing the probability that an event exceeds time t, is expressed as:

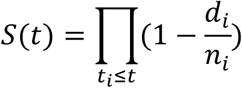

Where *t_i_* represents a time length of at least one event (e.g. one minute), *d_i_* the number of events at time *t_i_* (e.g. number of active times at one minute), and *n_i_* the number of events surviving (e.g. number of active times longer than one minute, either non-censored or right-censored) up to *t_i_*.

The python implementation ‘lifelines’ was used to calculate the Kaplan-Meier estimator^46^.

### Bootstrapped survival probabilities

Survival probabilities for burst size and burst amplitude were calculated from MS2 intensity traces. Only active transcriptional times starting and ending within the imaging window (uncensored) were considered. For resulting survival distributions, 95% confidence intervals were calculated from bootstrapping (n=10.000).

### Mathematical model and parameter fitting

The two-state, two-regime model was fitted to the data in two steps:

1. Single-Distance Fit: the model’s seven parameters were optimized using the data from a single enhancer-promoter distance (5 kb). The model was fitted simultaneously to the survival probabilities of the burst and inter-burst durations, the steady-state probability to observe a burst, and the correlation between consecutive inter-burst durations.
2. Multi-Distance Fit: for other enhancer-promoter distances, only one parameter was fitted while keeping the remaining parameters fixed at their 5 kb best-fit values. In this case, the model was only fitted to the survival probabilities of inter-burst durations.

The mathematical description and analysis of the model, and the fitting procedures are explained in detail in the **Supplementary Information**.

---

**Extended Data Figure 1.**
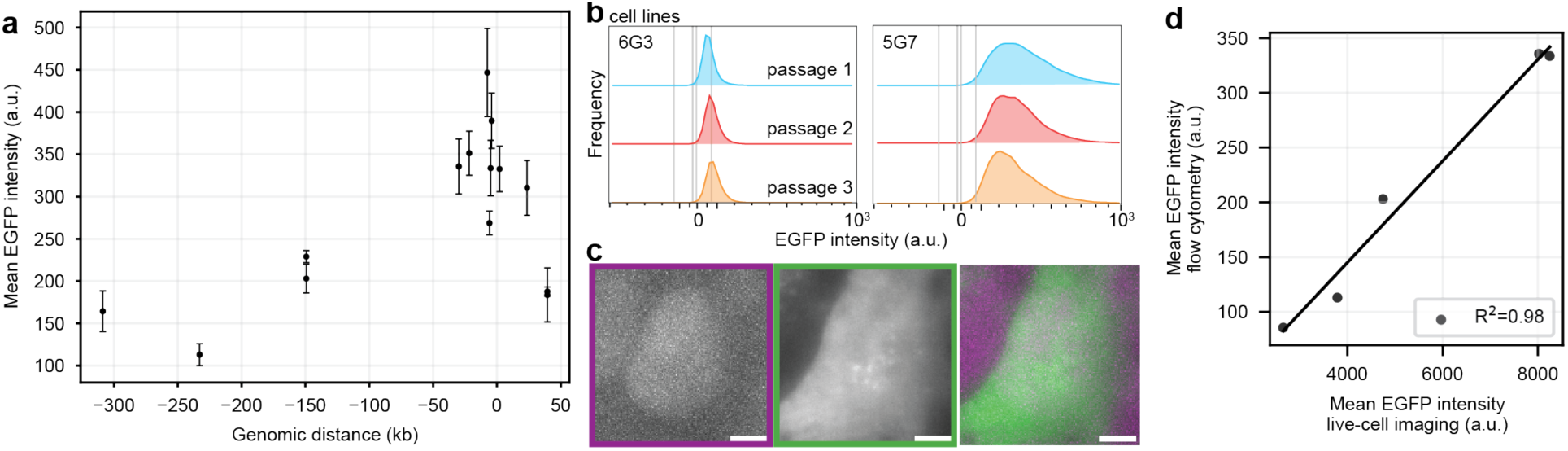
EGFP levels measured in live-cell imaging experiments and flow cytometry are correlated. **a.** EGFP levels of individual EGFP+ cell lines over cell passages measured by flow cytometry. **b.** Mean EGFP intensity assessed through flow cytometry in individual EGFP+ cell lines as a function of genomic distance between the promoter and the enhancer. **c.** Representative images from live-cell imaging of the MCP-HaloTag channel (640 nm laser, left), EGFP channel (480 nm laser, middle) and overlay (right). Scale bar: 5 µm. **d.** Pearson correlation of mean EGFP levels measured using flow cytometry with mean EGFP levels measured in live-cell imaging. Pearson coefficient of determination (r2) = 0.98.

**Extended Data Figure 2.**
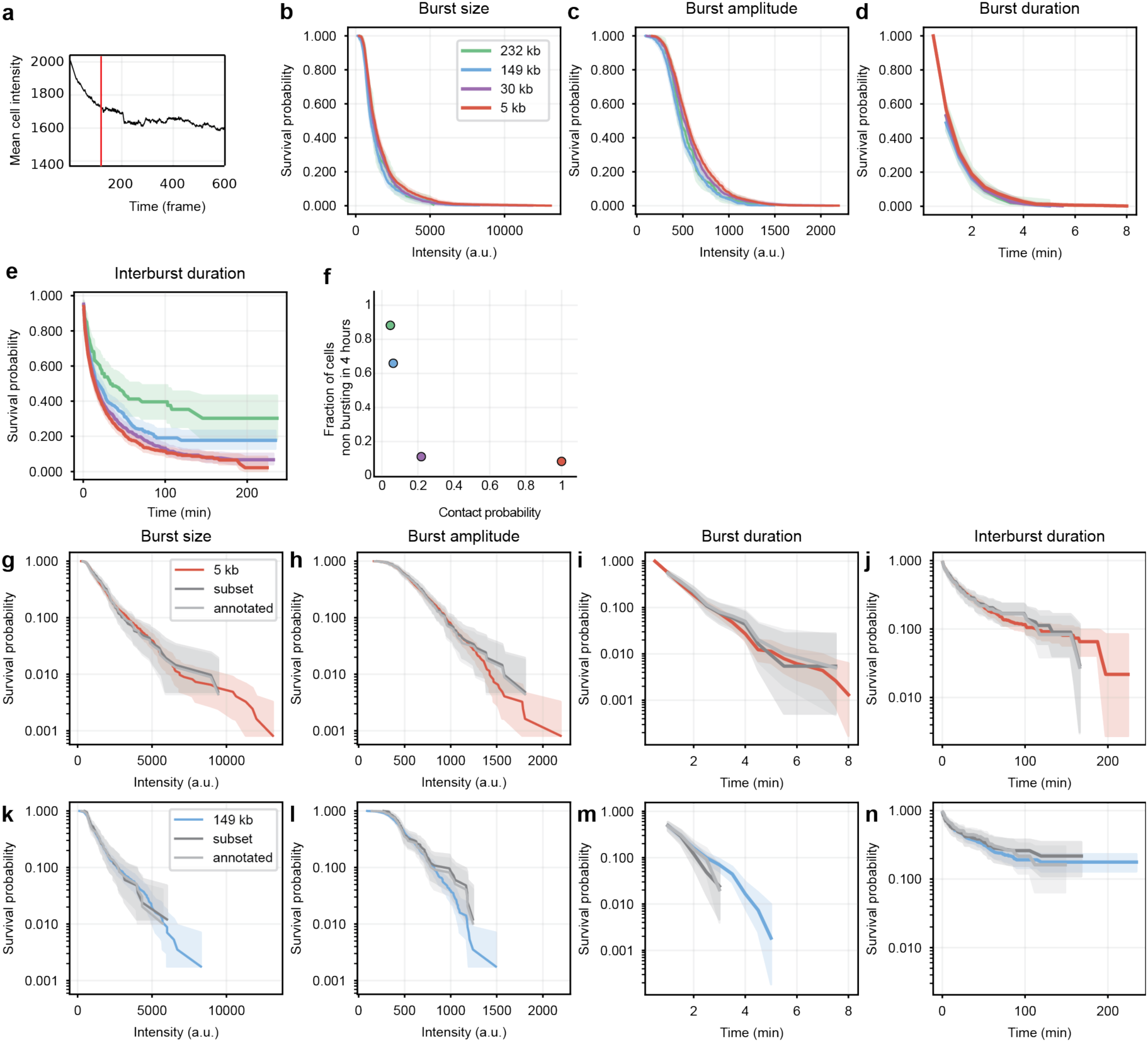
Quantification of burst parameters is representative of underlying dynamics. **a.** Average intensity of nuclear background as a function of frame number. Red line: lower bound (120 frames) the time window considered in the burst analysis. **(b)** Survival probability of burst size. 95% confidence intervals were estimated by bootstrapping (n=10.000). **(c)** Survival probability of burst amplitude. 95% confidence intervals were estimated by bootstrapping (n=10.000). **(d)** Survival probability of burst duration calculated with the Kaplan-Meier estimator. **(e)** Survival probability of inter-burst duration calculated with the Kaplan-Meier estimator. **f.** Fraction of non-bursting cells as function of contact probability as estimated by capture 3C^13^. **g-n.** comparison of burst parameters between the full dataset (colored line), a subset of the dataset (dark gray line) and manually annotated versions of the same subset (light gray line) for two clonal cell lines (upper panel = 5 kb enhancer-promoter genomic distance, lower panel = 149 kb enhancer-promoter genomic distance).

**Extended Data Figure 3.**
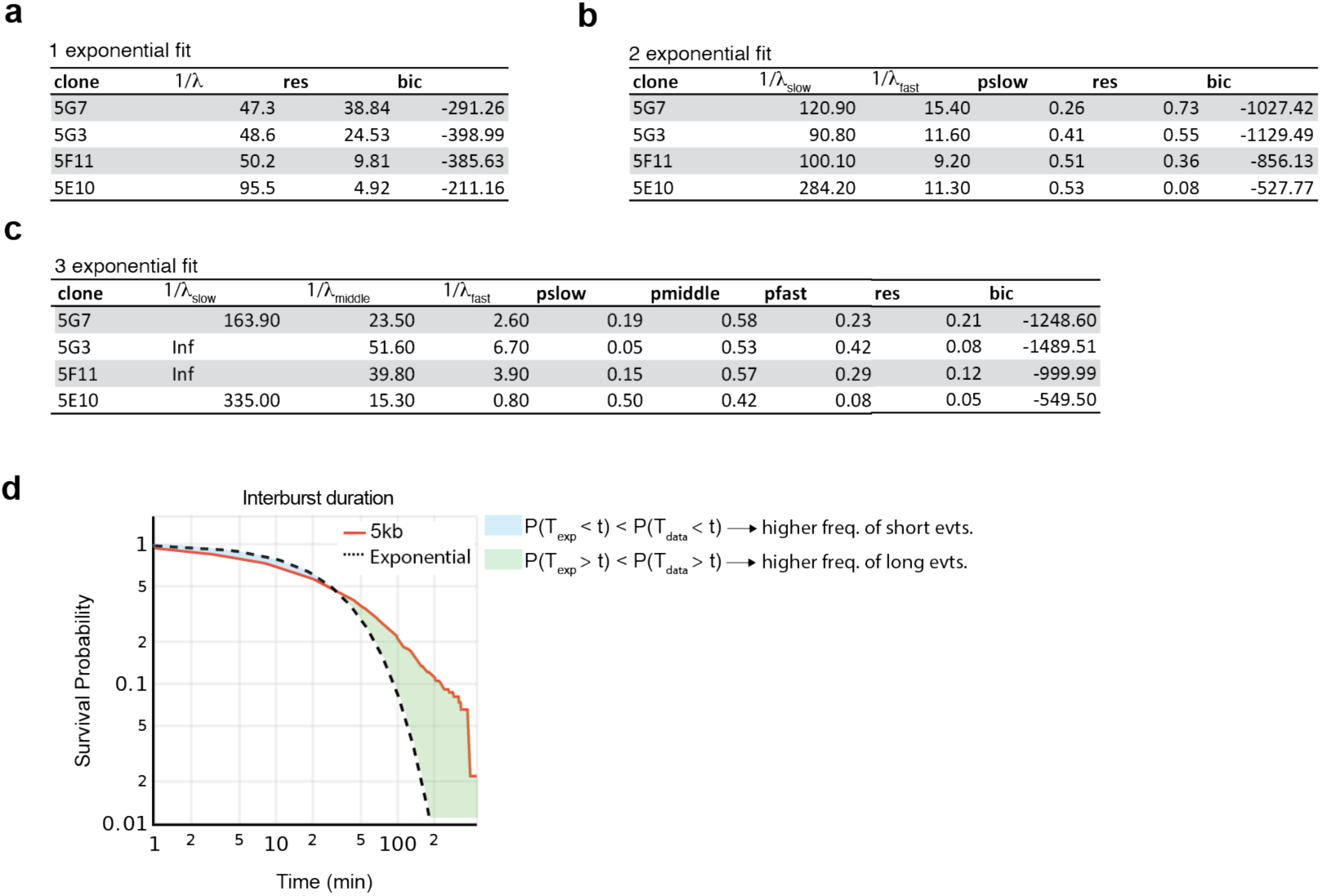
Inter-burst durations are distributed over three timescales. **a.** Parameter values for the best-fitting one exponential model to survival probabilities of inter-burst durations in cell lines where the SCR was located at 5 kb, 29 kb, 149 kb, and 232 kb from the promoter. The inverse of the best-fit exponent is shown in minutes. The residual of the least squares fit (res) and the Bayesian information criterion (bic) are shown. **b.** Similar to panel **(a),** but for a two-exponential model. Pslow is the weight associated with the slowest exponential. **c.** Similar to panel **(a),** but for a three-exponential model. **d.** Survival probabilities of the inter-burst durations for the cell line where the SCR was located at 5 kb from the promoter (red line). Survival probabilities of an exponential variable matching the median of the inter-burst durations (black dotted line). For a time t where the survival probabilities of the inter-burst durations is below (resp. above) the survival probabilities of the exponential, the probability that events lasted less (resp. more) than a time t is higher in the data than for a Poisson process (blue and green area, resp.).

**Extended Data Figure 4.**
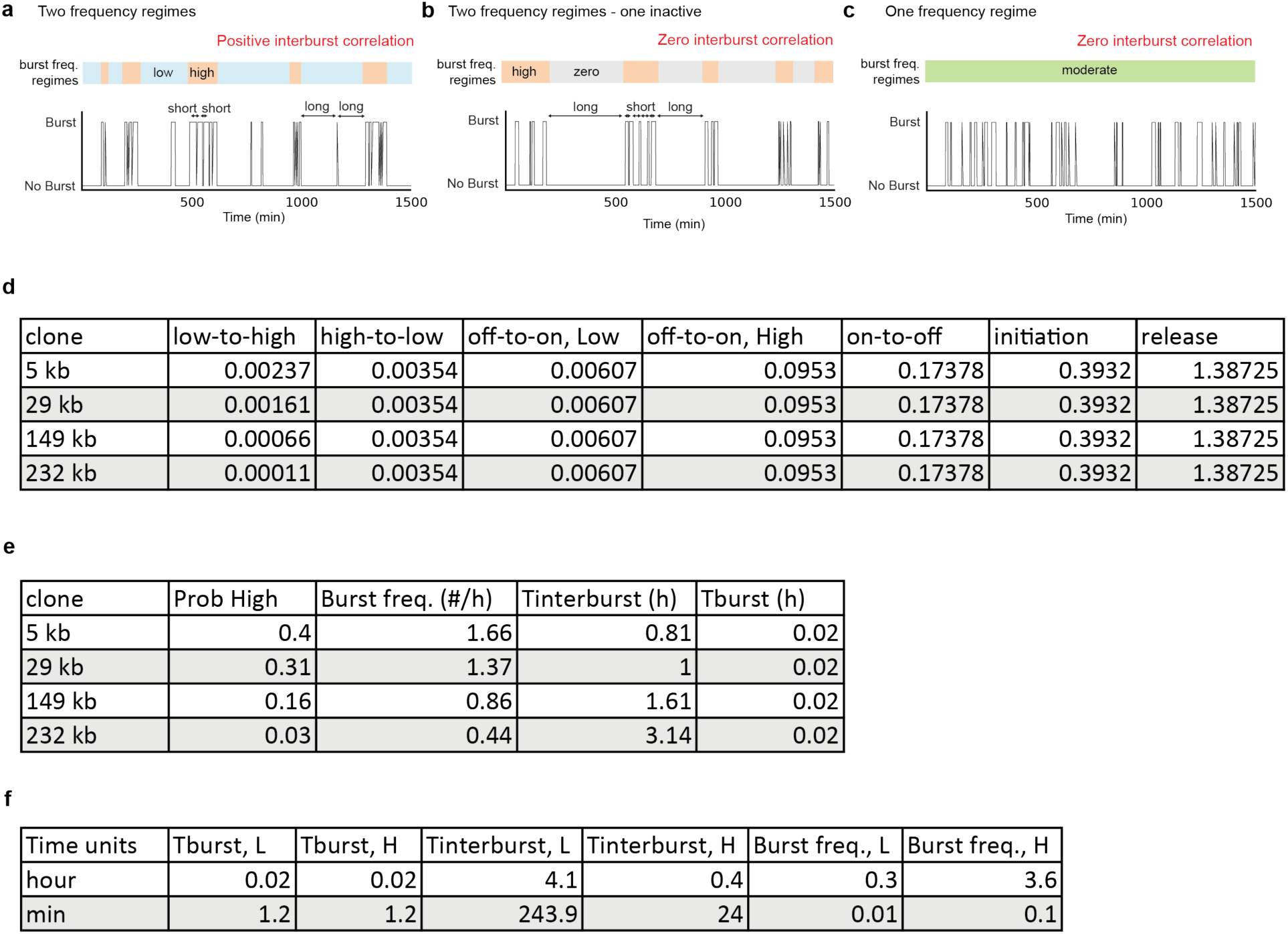
**a-c**. Schematic of the single-cell burst dynamics under the assumption of two frequency regimes (a), two frequency regimes with one of them being inactive (b), and one frequency regime (c). Sperman correlation between consecutive inter-burst periods can be positive when the promoter operates in two different active regimes (a) but is zero when only one regime is active (b-c). **d.** Parameter values for the best fitting two state, two regime model. All the parameters were fit to the data of the 5 kb clone, except for the low-to-high transition rate which was fit to each clone independently. The unit of the rates is 1/minute. **e**. Predictions of the best fitting model for the steady-state probability that the promoter is in the high-frequency regime, the burst frequency, the mean inter-burst duration, and the mean burst duration. **f**. Predictions of the best fitting model for mean burst and inter-burst duration and burst frequency in each of the regimes. Burst and inter-burst frequencies were calculated as the fraction of time spent in a burst (or inter-burst) divided by the mean duration of a burst (or inter-burst).

**Extended Data Figure 5.**
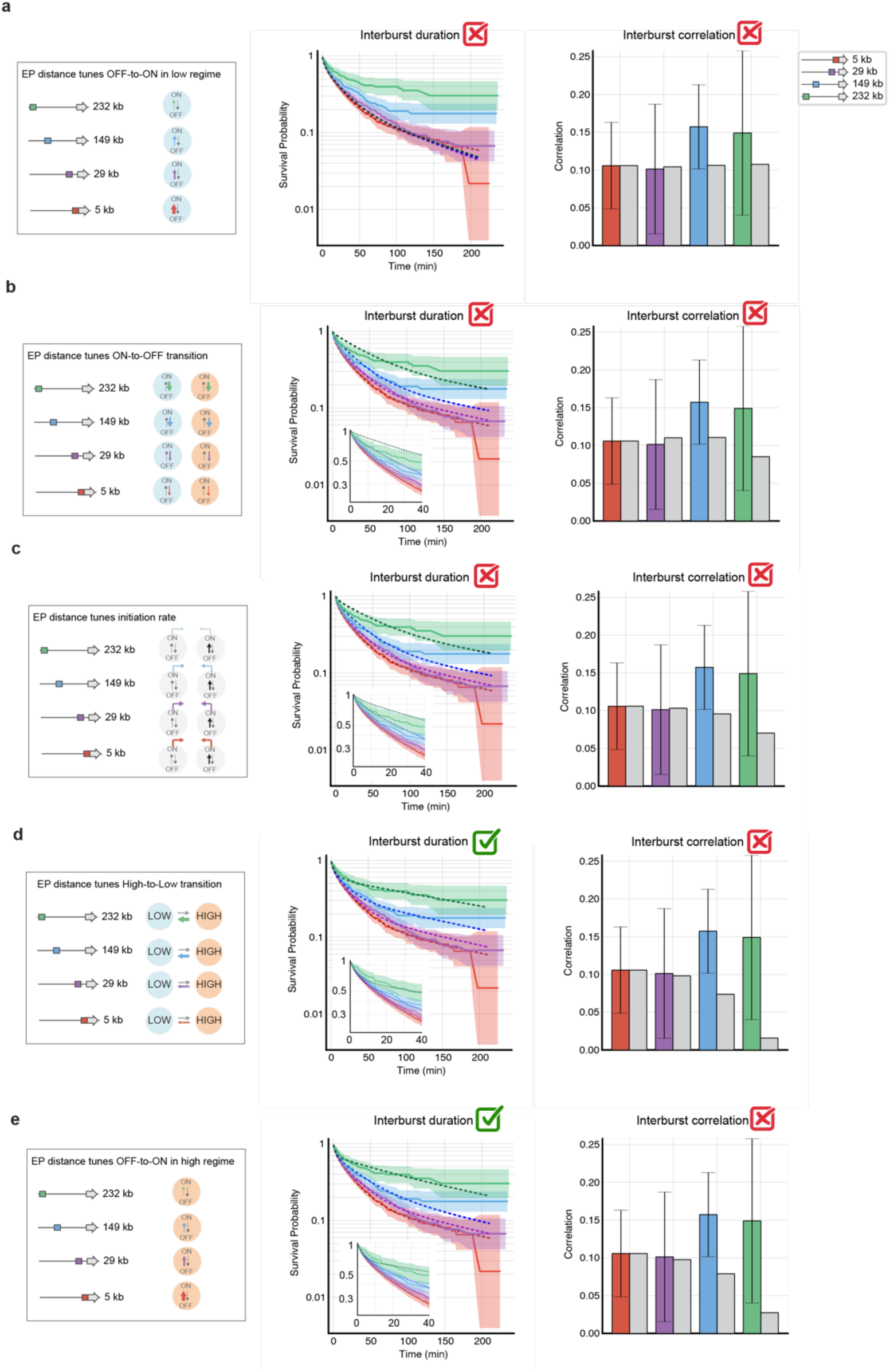
**a-e**. Best fit survival probabilities of the inter-burst duration of the 29 kb, 149 kb, and 232 kb cell lines (purple, blue, and green dotted lines) and predicted Pearson correlation of consecutive inter-burst periods, for models where the enhancer-promoter genomic distance only affects the OFF-to-ON rate in the low-frequency regime (a), the ON-to-OFF rate (b), the initiation rate (c), the high-to-low rate (d), and the OFF-to-ON rate in the high-frequency regime (e). Data is similar to Figure 6.

## Notes

### Competing Interest Statement

The authors have declared no competing interest.

