## Supplementary Information for "Enhancer control of promoter activity and variability *via* frequency modulation of clustered transcriptional bursts"

In this Supplementary Information, we develop a theoretical framework to analyze the bursting dynamics of promoters with arbitrary combinatorial complexity. This framework provides analytical expressions for key observables, including the survival probabilities of burst and inter-burst durations, the steady-state probability of observing a burst, and the correlation between consecutive inter-burst periods. We then apply this framework to analyze and fit the two-state, two-regime model introduced in the main text.

The reminder of this document is structured as follows. Section 1 introduces the class of stochastic multi-state transcription models. Sections 2 derives analytical expressions for (1) the survival probabilities of the burst and inter-burst durations, (2) the steady-state probability of observing a burst, and (3) the correlation between consecutive inter-burst periods. Section 3 extends these expressions to account for the removal of singlet bursts from the data. Finally Sections 4 and 5 present the two-state, two-regime model and describe our method for fitting it to experimental data.

#### 1 Generic stochastic multi-state model of transcription

We consider a generic multi-state transcription model where a promoter can occupy one of $n$ states, denoted $\{s_1, \dots, s_n\}$, and undergo stochastic transitions between them at rates $k_{ij}$. From state $s_i$, transcription is initiated stochastically at rate $\mu_i$. States with $\mu_i = 0$ are "OFF" states, and states with $\mu_i > 0$ are "ON" states. Upon initiation, an mRNA stem loop reporter forms instantaneously and is stochastically released from DNA at a rate $\delta$. The kinetic reactions governing this model are:

$$\begin{aligned}
s_i &\underset{k_{ji}}{\overset{k_{ij}}{\rightleftharpoons}} s_j, \quad \forall i \neq j, \quad i, j \in \{1, \dots, n\} \\
s_i &\underset{\mu_i}{\rightarrow} s_i + \text{mRNA}, \quad \forall i \in \{1, \dots, n\} \\
\text{mRNA} &\underset{\delta}{\rightarrow} \emptyset
\end{aligned}$$

This system defines a continuous-time Markov chain, $Z_t = (z_1(t), z_2(t))$, where $z_1(t)$ represents the promoter state at time $t$, and $z_2(t)$ represents the number of nascent mRNA at time $t$.

#### 1.1 Linear indexing of the state space

To facilitate computations, we introduce a *linear indexing function* mapping the pair $(z_1, z_2)$ to a single "hyper" state $\phi(z_1, z_2) \in \mathbb{N}$, defined as:

$$\phi(i, m) = n(m - 1) + i, \tag{1}$$

with its inverse

$$\phi^{-1}(k) = (k \bmod n, \lfloor \frac{k}{n} \rfloor). \tag{2}$$

In other words, this indexing orders the state space such that the first $n$ states correspond to promoter states with $0$ mRNA, the next $n$ states correspond to promoter states with $1$ mRNA, and so forth. The following scheme illustrates the linear indexing for $5$ promoter states:

$s_1 \quad s_2 \quad s_3 \quad s_4 \quad s_5 \quad s_1 \quad s_2 \quad s_3 \quad s_4 \quad s_5 \quad s_1 \quad s_2 \quad s_3 \quad s_4 \quad s_5 \quad \cdots$

mRNA = 0 mRNA = 1 mRNA = 2

#### 1.2 Infinitesimal generator of the process $Z_t$

The infinitesimal generator of the continuous-time Markov chain $Z_t$ is the matrix $\mathbf{Q}$ containing all transition rates, with each row summing to zero. Based on the definition of the multi-state transcription model, the generator decomposes as

$$\mathbf{Q} := \mathbf{Q}_p + \mathbf{T} + \mathbf{D} \tag{3}$$

where:

- $\mathbf{Q}_p$ governs promoter state transitions for each number of nascent mRNA,
- $\mathbf{T}$ describes transcription initiation,
- $\mathbf{D}$ accounts for mRNA release.

The matrix $\mathbf{Q}_p$ describes transitions between promoter states while maintaining the same number of nascent mRNA:

$$\mathbf{Q}_p = \mathbf{I} \otimes \mathbf{K} \tag{4}$$

where $\mathbf{I}$ is the identity matrix, $\mathbf{K} = (k_{ij})$ is the $n \times n$ promoter transition matrix, and $\otimes$ is the Kronecker product.

The matrix $\mathbf{T}$ describes the transition from $m$ to $m + 1$ mRNA:

$$\mathbf{t}_{i,j} = \begin{cases} \mu_{\{i \bmod n\}}, & \text{if } j = i + n \\ 0, & \text{otherwise} \end{cases} \tag{5}$$

The matrix $\mathbf{D}$ describes the transition from $m$ to $m - 1$ mRNA:

$$\mathbf{d}_{i,j} = \begin{cases} \lfloor i/n \rfloor \delta, & \text{if } j = i - m \\ 0, & \text{otherwise} \end{cases} \tag{6}$$

#### 1.3 Finite-state approximation

The state space of $Z_t$ is infinite because the number of nascent mRNA is unbounded. However, in practice, this number remains small. Inspired by the finite state projection method (Munsky and Khammash [2006]), we approximate the system by truncating the state space. Effectively, we define a maximal number of nascent mRNA per cell, $M_{\text{max}}$, leading to a finite state space:

$$\tilde{Z}_t \in \{1, \ldots, n\} \times \{0, \ldots, M_{\text{max}}\}. \tag{7}$$

The infinitesimal generator of the truncated process, $\tilde{\mathbf{Q}}$, has entries:

$$\tilde{q}_{i,j} = \begin{cases} q_{ij}, & \text{if } i \neq j \\ -\sum_{k \leq n(M_{\text{max}}+1), k \neq i} q_{i,k} & \text{if } i = j \end{cases} \tag{8}$$

#### 1.4 Discretization of the generic model

Experimental observations are acquired every 30 seconds. To match this time resolution, we define the discretized process:

$$X_k := \tilde{Z}_{k\Delta t}, \tag{9}$$

where $\Delta t = 30s$. The process $X_k$ forms a discrete-time Markov chain in $\{1, \ldots, n\} \times \{0, \ldots, M_{\text{max}}\}$, with transition matrix $\mathbf{P} = (p_{i,j})$, computed directly from the infinitesimal generator $\tilde{\mathbf{Q}}$ via:

$$p_{ij} = \mathbb{P}(X_{k+1} = j | X_k = i) = (e^{\Delta t \tilde{\mathbf{Q}}})_{ij}. \tag{10}$$

The steady-state distribution $\boldsymbol{\pi}$ of the Markov chain $X$ is the left eigenvector of the transition matrix $\mathbf{P}$ associated with the eigenvalue 1.

#### 1.5 What is a *burst* in the model?

In our experiments, the final readout is a binary track for each cell, where 1 and 0 correspond to the presence or absence of nascent mRNA at the gene, respectively. A ”burst” is defined as a period of consecutive ones. In the model, we define a similar binary process $B_t$, where

- $B_t = 1$ if there is at least one nascent mRNA at the promoter,
- $B_t = 0$ ohtherwise

Using the linear indexing of the state space, we define the set of non-bursting states:

$$\mathcal{N} = \{1, \ldots, n\}, \tag{11}$$

and the set of ”bursting” states:

$$\mathcal{B} = \{n + 1, \ldots, n(M_{max} + 1)\}. \tag{12}$$

Thus, the burst process is given by:

$$B_t = \begin{cases} 1, & \text{if } X_t \in \mathcal{B} \\ 0 & \text{if } X_t \in \mathcal{N}. \end{cases} \tag{13}$$

### 2 Computation of the observables in the generic model

In this section we derive matrix-based formulas to compute the survival probabilities of burst and inter-burst durations, the steady-state probability of observing a burst, and the correlation between consecutive inter-burst durations. These calculations ultimately reduce to computing the eigenvector of $\mathbf{P}$ corresponding to eigenvalue $1$ (steady-state distribution), inverting matrices of the form $\mathbf{I} - \mathbf{P}_{\mathcal{A}}$ for sub-matrices $\mathbf{P}_{\mathcal{A}}$ of the matrix $\mathbf{P}$, and performing direct matrix multiplications.

#### 2.1 Survival probabilities of the burst durations

To compute the distribution of burst durations, we define a new absorbing Markov chain $X^{\mathcal{B}}$, which modifies the original process $X$ by making all non-bursting states ($\mathcal{N}$) absorbing. In other words, the modified process $X^{\mathcal{B}}$ evolves like the original process $X$ until it reaches a state with zero nascent mRNA, at which point it stops permanently. The transition matrix of the absorbing Markov chain $X^{\mathcal{B}}$ is:

$$\tilde{\mathbf{P}} = \begin{bmatrix} \mathbf{P}_{\mathcal{BB}} & \mathbf{P}_{\mathcal{BN}} \\ \mathbf{1} & \mathbf{0} \end{bmatrix} \tag{14}$$

where $\mathbf{P}_{\mathcal{AB}}$ represents the transitions between states in $\mathcal{A}$ and $\mathcal{B}$.

The probability that a burst, starting from state $i$ in the original process $X$, lasts longer than time $t$ is equivalent to the probability that a burst, starting from $i$ in the modified process $X^{\mathcal{B}}$, has not yet been absorbed at time $t$:

$$\mathbb{P}(T_{\text{Burst}} > t | X_0^{\mathcal{B}} = i) = \sum_{j\in\mathcal{B}} (\mathbf{P}_{\mathcal{BB}}^t)_{ij}. \tag{15}$$

To compute the steady-state probability of a burst lasting more than $t$, we need to determine the steady-state distribution of states at which bursts start, denoted by $\mathbf{p}^{\mathcal{B}}$:

$$\begin{aligned} p_j^{\mathcal{B}} &:= \mathbb{P}(\text{Burst starts in state } j) \\ &= \mathbb{P}(X_{t+1} = j | X_t \in \mathcal{N}, X_{t+1} \in \mathcal{B}) \\ &= \frac{\mathbb{P}(X_{t+1} = j, X_t \in \mathcal{N})}{\mathbb{P}(X_{t+1} \in \mathcal{B}, X_t \in \mathcal{N})} \end{aligned} \tag{16}$$

We expend the numerator using the Markov property:

$$\begin{aligned} \mathbb{P}(X_{t+1} = j, X_t \in \mathcal{N}) &= \sum_{i\in\mathcal{N}} \mathbb{P}(X_{t+1} = j, X_t = i) \\ &= \sum_{i\in\mathcal{N}} \mathbb{P}(X_{t+1} = j | X_t = i)\mathbb{P}(X_t = i) \end{aligned}$$

Expressed in terms of the steady-state probability distribution $\boldsymbol{\pi}$ of $X$ and its transition matrix $\mathbf{P}$,

$$\mathbb{P}(X_{t+1} = j, X_t \in \mathcal{N}) = \sum_{i\in\mathcal{N}} p_{ij}\pi_i \tag{17}$$

Substituting equation (17) in equation (16), we obtain

$$\mathbf{p}^{\mathcal{B}} = \frac{1}{\boldsymbol{\pi}^{\mathcal{N}}\mathbf{P}_{\mathcal{NB}}\mathbf{e}} \boldsymbol{\pi}^{\mathcal{N}}\mathbf{P}_{\mathcal{NB}}$$

where $\boldsymbol{\pi}^{\mathcal{N}}$ is the row vector with entries describing the steady-state probabilities of the non-bursting states, and $\mathbf{e}$ is the column vector with all entries equal to 1.

Combining the previous results, the probability that a burst lasts for more than time $t$ is given by:

$$\mathbb{P}(T_{\text{Burst}} > t) = \mathbf{p}^{\mathcal{B}} \mathbf{P}_{\mathcal{B}\mathcal{B}}^{t} \mathbf{e}. \tag{18}$$

#### 2.2 Survival probabilities of the inter-burst durations

The survival probability of inter-burst durations is computed using the same strategy as for the burst durations, but with the bursting states $\mathcal{B}$ as absorbing. First, we define the steady-state distribution of the states where an inter-burst period begins:

$$\mathbf{p}^{\mathcal{N}} = \frac{1}{\boldsymbol{\pi}^{\mathcal{B}} \mathbf{P}_{\mathcal{B}\mathcal{N}} \mathbf{e}} \boldsymbol{\pi}^{\mathcal{B}} \mathbf{P}_{\mathcal{B}\mathcal{N}} \tag{19}$$

where $\boldsymbol{\pi}^{\mathcal{B}}$ is the row vector with entries describing the steady-state probabilities of the bursting states, and $\mathbf{e}$ is the column vector with all entries equal to 1. The survival probability of an inter-burst durations is then:

$$\mathbb{P}(T_{\text{Inter-burst}} > t) = \mathbf{p}^{\mathcal{N}} \mathbf{P}_{\mathcal{N}\mathcal{N}}^{t} \mathbf{e}. \tag{20}$$

#### 2.3 Probability to observe a burst

The steady state probability to observe a burst is given by the probability that the Markov chain $X$ is in a bursting state at steady-state:

$$\begin{aligned} \mathbb{P}(\text{to observe a burst}) &= \mathbb{P}(X_t \in \mathcal{B}) \\ &= \sum_{i \in \mathcal{B}} \pi_i \end{aligned}$$

#### 2.4 Correlation between consecutive inter-burst periods

Let $T_1$ and $T_2$ denote the durations of two consecutive inter-burst periods. Their correlation is defined as:

$$\rho_{T_1,T_2} := \frac{\text{Cov}(T_1, T_2)}{\sigma_{T_1} \sigma_{T_2}} \tag{21}$$

where $\text{Cov}(T_1, T_2) = \mathbb{E}[(T_1 - \mathbb{E}[T_1])(T_2 - \mathbb{E}[T_2])]$ is the covariance between the two durations, and $\sigma_T = \sqrt{\text{Var}(T)}$ is the standard deviation of $T$. Since $T_1$ and $T_2$ are identically distributed, we can express the covariance as:

$$\text{Cov}(T_1, T_2) = \mathbb{E}[T_1 T_2] - \mathbb{E}[T_1]^2, \tag{22}$$

and the denominator simplifies to:

$$\sigma_{T_1} \sigma_{T_2} = \text{Var}(T_1) = \mathbb{E}[T_1^2] - \mathbb{E}[T_1]^2.$$

To compute the expectations $\mathbb{E}[T_1^2]$, $\mathbb{E}[T_1]$, $\mathbb{E}[T_1 T_2]$, we use the absorbing Markov chains introduced in Sections 2.1 and 2.2:

- $X^{\mathcal{N}}$, which evolves like the original process $X$ until it reaches a bursting state, where it is absorbed.

- $X^{\mathcal{B}}$, which evolves like $X$ until it reaches a non-bursting state, where it is absorbed.

The inter-burst duration $T_1$ is equivalent to the time before absorption in the Markov chain $X^{\mathcal{N}}$. A key element for computing absorbing times is the fundamental matrix,

$$\mathbf{N}^{\mathcal{N}} = (\mathbf{I} - \mathbf{P}_{\mathcal{N}\mathcal{N}})^{-1}. \tag{23}$$

The $(i-j)$-th entry of $\mathbf{N}^{\mathcal{N}}$ represents the expected time spent in state $j$ given that $X^{\mathcal{N}}$ started in state $i$. Similarly, the fundamental matrix for $X^{\mathcal{B}}$ is:

$$\mathbf{N}^{\mathcal{B}} = (\mathbf{I} - \mathbf{P}_{\mathcal{B}\mathcal{B}})^{-1}. \tag{24}$$

Following Iosifescu [1980], the first and second moments of the absorbing time are:

$$\mathbb{E}[T_i] \;=\; \mathbf{p}^{\mathcal{N}}\mathbf{N}^{\mathcal{N}}\mathbf{1} \tag{25}$$

$$\mathbb{E}[T_i^2] \;=\; \mathbf{p}^{\mathcal{N}}\left((2\mathbf{N}^{\mathcal{N}} - \mathbf{I})\mathbf{N}^{\mathcal{N}}\mathbf{1}\right) \tag{26}$$

$$\tag{27}$$

for $i = 1, 2$, where $\mathbf{p}^{N}$ is the steady-state probability of the initial state of an inter-burst event, as defined in equation (19).

Unlike $\mathbb{E}[T_1]$ and $\mathbb{E}[T_1^2]$, the expectation $\mathbb{E}[T_1T_2]$ cannot be directly obtained from the absorbing Markov chain $X^{\mathcal{N}}$, since it depends on the burst separating the two inter-burst periods. Using conditional probability, we expand:

$$E[T_1T_2] = \sum_{t\geq 0} t\mathbb{E}[T_2|T_1 = t]\mathbb{P}(T_1 = t). \tag{28}$$

By conditioning on the final state of the first inter-burst period:

$$\mathbb{E}[T_2 \mid T_1 = t] = \sum_{j\in\mathcal{B}} \mathbb{E}[T_2 \mid X_{T_1} = j]\,\mathbb{P}(X_{T_1} = j \mid T_1 = t). \tag{29}$$

Further conditioning on the state at the end of the burst separating $T_1$ and $T_2$:

$$\mathbb{E}[T_2 \mid X_{T_1} = j] = \sum_{k\in\mathcal{N}} \mathbb{E}[T_2 \mid X_\tau = k]\,\mathbb{P}(X_\tau = k \mid X_{T_1} = j), \tag{30}$$

where $\tau$ is the burst duration separating the two inter-burst events. Substituting, we obtain:

$$\mathbb{E}[T_1T_2] = \sum_{t\geq 0}\sum_{j\in\mathcal{B}}\sum_{k\in\mathcal{N}} t\mathbb{E}[T_2 \mid X_\tau = k]\,\mathbb{P}(X_\tau = k \mid X_{T_1} = j)\mathbb{P}(X_{T_1} = j \mid T_1 = t)\mathbb{P}(T_1 = t). \tag{31}$$

The second term in equation (31) is the expectation of the inter-burst duration, given the initial state $k$, which correspond to the time to absorption of the Markov chain $X^{\mathcal{N}}$ given its initial state:

$$\mathbb{E}[T_2|X_\tau = j] = (\mathbf{N}^{\mathcal{N}}\mathbf{e})_j. \tag{32}$$

The third term in equation (31) is the probability that the Markov chain $X^{\mathcal{B}}$ is absorbed in the non-bursting state $k$, given that it starts in the bursting state $j$. Following Iosifescu [1980], this probability corresponds to the $j, k$ entry of the matrix $\mathbf{N}^{\mathcal{B}}\mathbf{P}_{\mathcal{B}\mathcal{N}}$:

$$\mathbb{P}(X_\tau = k|X_{T_1} = j) = (\mathbf{N}^{\mathcal{B}}\mathbf{P}_{\mathcal{B}\mathcal{N}})_{jk}. \tag{33}$$

The fourth term in equation (31) is the distribution of states at the end of the first inter-burst duration, conditioned on the duration of the inter-burst:

$$\begin{aligned}
\mathbb{P}(X_{T_1} = j | T_1 = t) &= \mathbb{P}(X_t = j | T_1 = t) \\
&= \frac{\mathbb{P}(X_t = j, T_1 = t)}{\mathbb{P}(T_1 = t)} \\
&= \frac{1}{\mathbb{P}(T_1 = t)} \sum_{\ell \in \mathcal{N}} \mathbb{P}(X_t = j, X_{t-1} = \ell) \\
&= \frac{1}{\mathbb{P}(T_1 = t)} \sum_{\ell \in \mathcal{N}} \mathbb{P}(X_t = j | X_{t-1} = \ell) \mathbb{P}(X_{t-1} = \ell) \\
&= \frac{1}{\mathbb{P}(T_1 = t)} \sum_{\ell \in \mathcal{N}} \sum_{q \in \mathcal{N}} \mathbb{P}(X_t = j | X_{t-1} = \ell) \mathbb{P}(X_{t-1} = \ell | X_0 = q) \mathbb{P}(X_0 = q) \\
&= \frac{1}{\mathbb{P}(T_1 = t)} \sum_{\ell \in \mathcal{N}} (\mathbf{P}_{\mathcal{N}\mathcal{B}})_{\ell j} (\mathbf{P}_{\mathcal{N}\mathcal{N}}^{t-1})_{q\ell} \mathbf{p}_q^{\mathcal{N}} \qquad (34)
\end{aligned}$$

Substituting equations (32), (33), and (34) into equation (31), we obtain the final matrix formulation:

$$E[T_1 T_2] = \sum_{t \geq 0} t \mathbf{p}^{\mathcal{N}} \mathbf{P}_{\mathcal{N}\mathcal{N}}^{t-1} \mathbf{P}_{\mathcal{N}\mathcal{B}} N_{\mathcal{B}} \mathbf{P}_{\mathcal{B}\mathcal{N}} \mathbf{N}_{\mathcal{N}} \mathbf{e} \qquad (35)$$

### 3 Computation of the observables after removing singlet bursts

In the final bursting track obtained for each cell, singlet bursts (i.e., bursts lasting only one frame) were filtered out. To ensure a consistent comparison between the experimental data and the model, we must apply the same filtering to the model-generated bursts. For burst durations, this is straightforward: we simply condition on bursts lasting more than one frame. For inter-burst durations, filtering is more complex because removing a singlet burst merges two consecutive inter-burst periods, altering their distribution.

To systematically remove singlet bursts in the process $X$, we expand the state space by introducing a set of *pre-bursting* states, $\mathcal{B}^p$, which are copies of the bursting states. The new state space consists of:

$$\begin{aligned}
\mathcal{N} &= \{1, \ldots, n\} \\
\mathcal{B}^p &= \{n+1, \ldots, n(M_{max}+1)\} \\
\mathcal{B} &= \{n(M_{max}+1)+1, \ldots, 1n(M_{max}+1\}.
\end{aligned}$$

The key idea is to redirect the original process $X$ into $\mathcal{B}^p$ whenever it enters a bursting state, and then return it to the original state space in the next time step. In this way, we can discard the single visit in the pre-bursting states which correspond to the singlet bursts of the original process. Formerly, we define a modified process, $Y_t$, which evolves like $X_t$, except when entering a bursting state. Specifically, if $X_t$ enters the bursting state $i$ at time $t$ (i.e. $X_t = i$), the modified process $Y_t$ transitions into the corresponding pre-bursting state in $\mathcal{B}^p$ (i.e. $Y_t = i + nM_{max}$). In the next time step, the process $Y$ goes back to the original state space by following the same transition as $X$, i.e., $Y_{t+1} = X_{t+1}$.

We define the new binary process, $\tilde{B}_t$, that excludes singlet bursts as:

$$\tilde{B}_t = \begin{cases} 1, & \text{if } Y_t \in \mathcal{B} \\ 0 & \text{if } Y_t \in \mathcal{N} \cup \mathcal{B}^p \end{cases} \qquad (36)$$

The trajectories of $\tilde{B}_t$ match those of $B_t$ after singlet bursts are removed, except that bursts in $\tilde{B}_t$ are one time unit shorter because time spent in $\mathcal{B}^p$ is unrecorded. Thus, the probability that the inter-burst duration in $B_t$ (with singlet bursts removed) exceeds $t$ is equivalent to the probability that the inter-burst duration in $\tilde{B}_t$ exceeds $t+1$:

$$\mathbb{P}(T_{\text{Inter-burst, w/o singlets}} > t) = \mathbb{P}(\tilde{T}_{\text{Inter-burst}} > t+1) \tag{37}$$

The transition matrix $\tilde{\mathbf{P}}$ of the process $Y$ is structured as:

$$\tilde{\mathbf{P}} = \begin{array}{c} \begin{array}{ccc} \mathcal{N} & \mathcal{B}^p & \mathcal{B} \end{array} \\ \left[\begin{array}{c|c|c} \mathbf{P}_{\mathcal{N}\mathcal{N}} & \mathbf{P}_{\mathcal{N}\mathcal{B}} & \mathbf{0} \\ \hline \mathbf{P}_{\mathcal{B}\mathcal{N}} & \mathbf{0} & \mathbf{P}_{\mathcal{B}\mathcal{B}} \\ \hline P_{\mathcal{B}\mathcal{N}} & \mathbf{0} & \mathbf{P}_{\mathcal{B}\mathcal{B}} \end{array}\right] \end{array} \begin{array}{c} \mathcal{N} \\ \mathcal{B}^p \\ \mathcal{B} \end{array} \tag{38}$$

To simplify notation, wemerge the original non bursting states $\mathcal{N}$ and pre-bursting states $\mathcal{B}^p$ into a single set $\tilde{\mathcal{N}}$. The transition matrix then simplifies to :

$$\tilde{\mathbf{P}} = \begin{bmatrix} \mathbf{P}_{\tilde{\mathcal{N}}\tilde{\mathcal{N}}} & \mathbf{P}_{\tilde{\mathcal{N}}\mathcal{B}} \\ \mathbf{P}_{\mathcal{B}\tilde{\mathcal{N}}} & \mathbf{P}_{\mathcal{B}\mathcal{B}} \end{bmatrix} \tag{39}$$

#### 3.1 Survival probabilities of the inter-burst durations

To compute the survival probability of the inter-burst duration after removing the singlet bursts, we apply the strategy outlined in Section 2.1 to the Markov chain $Y$, treating bursting states ($\mathcal{B}$) as absorbing. From equation (37), we obtain

$$\mathbb{P}(T_{\text{Inter-burst, wo singlets}} > t) = \mathbf{p}^{\tilde{\mathcal{N}}} \mathbf{P}_{\tilde{\mathcal{N}}\tilde{\mathcal{N}}}^{t+1} \mathbf{e}, \tag{40}$$

where $\mathbf{p}^{\tilde{\mathcal{N}}}$ is the steady-state distribution of states where an inter-burst period starts in the Markov chain $Y$.

#### 3.2 Correlation between consecutive inter-burst durations

To compute the correlation between consecutive inter-burst durations after removing the singlet bursts, we apply the strategy outlined in Section 2.4 to the Markov chain $Y$. Let

- $T_1^w$ and $T_2^w$ be the durations of two consecutive inter-burst periods in the original Markov chain $X$ after singlet bursts are removed.
- $\tilde{T}_1$ and $\tilde{T}_2$ be the corresponding durations in the Markov chain $Y$.

Since $T_i^w = \tilde{T}_i + 1$, the correlation between $T_1^w$ and $T_2^w$ is equivalent to the one between $\tilde{T}_1$ and $\tilde{T}_2$, leading to:

$$\rho_{T_1^w, T_2^w} = \frac{\mathbb{E}[\tilde{T}_1\tilde{T}_2] - \mathbb{E}[\tilde{T}_1]^2}{\mathbb{E}[\tilde{T}_1^2] - \mathbb{E}[\tilde{T}_1]^2}. \tag{41}$$

Following equations (26), (27) and (35), we compute each term in equation (41) as follows:

$$\mathbb{E}[T_i^w] = \mathbf{p}^{\tilde{\mathcal{N}}} \tilde{\mathbf{N}} \mathbf{1} \tag{42}$$

$$\mathbb{E}[(T_i^w)^2] = \mathbf{p}^{\tilde{\mathcal{N}}} \left( (2\tilde{\mathbf{N}}^{\tilde{\mathcal{N}}} - \mathbf{I}) \tilde{\mathbf{N}}^{\tilde{\mathcal{N}}} \mathbf{1} \right) \tag{43}$$

$$E[T_1^w T_2^w] = \sum_{t \geq 0} t \mathbf{p}^{\tilde{\mathcal{N}}} \mathbf{P}_{\tilde{\mathcal{N}}\tilde{\mathcal{N}}}^{t-1} \mathbf{P}_{\tilde{\mathcal{N}}\mathcal{B}} \mathbf{N}_{\mathcal{B}} \mathbf{P}_{\mathcal{B}\tilde{\mathcal{N}}} \tilde{\mathbf{N}}_{\tilde{\mathcal{N}}} \mathbf{e} \tag{44}$$

### 4 Two-state, two-regime model

The two-state, two-regime model is a stochastic, multi-state transcription model where the promoter can switch between two different transcriptional regimes: 'low-frequency' and 'high-frequency' regimes. In each regime, the promoter has an ON state, where transcription initiates at rate $\mu$, and an OFF state, where transcription is inactive. The only difference between the two regimes is the OFF-to-ON transition rates: $k_o n^L$ in the low-frequency regime and $k_{on}^H$ in the high-frequency, where $k_{on}^L < k_{on}^H$. In both regimes, the ON-to-OFF transition rate is $k_{off}$, and each RNA is released from DNA at rate $\delta$. The promoter can switch between regimes at rates $k_f$ from low- to high-frequency regime, and $k_b$ from high- to the low-frequency regime. This model consists of four states (two ON and two OFF), with the following kinetic scheme:

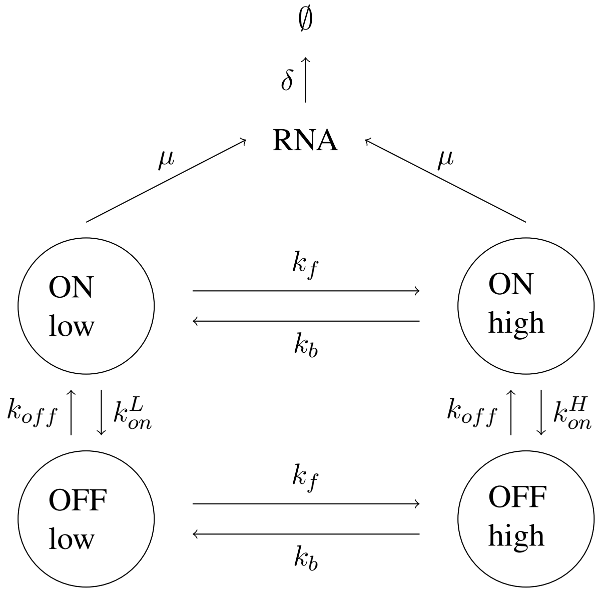


### 5 Model fitting

The two-state, two-regime model was fitted to live-cell imaging data in two steps:

1. *Single-Distance Fit*: The model's seven parameters were optimized using data from a single enhancer-promoter distance (5 kb).
2. *Multi-Distance Fit*: For other enhancer-promoter distances, only one parameter was fitted while keeping the remaining parameters fixed at their 5 kb best-fit values.

The following subsections describe the fitting strategy in detail.

#### 5.1 Fitting to a single distance

For the 5 kb enhancer-promoter distance, we fitted simultanuously the model to four experimental observables: (1-2) the survival probabilities of the burst and inter-burst durations, (3) the steady-state probability of observing a burst, and (4) the correlation between consecutive inter-burst durations.

The fitting minimized the weighted sum of squared errors, normalized by the number of data points. To account for both linear and logarithmic scaling in survival probabilities, we minimized a combination of squared errors and squared log errors.

For the optimization, we used a global search approach followed by a local refinement. For the global search, we used the *SAMIN* method (Simulated Annealing) from the Julia package *OptimizationOptimJL*. For local search, we used the *BFGS* method (Broyden–Fletcher–Goldfarb–Shanno algorithm) from the same package.

#### 5.2 Fitting across all distances

For each enhancer-promoter distance (except 5 kb), we fitted the model only to the survival probabilities of the inter-burst durations. Each fit optimized one free parameter at a time, while the remaining six parameters were fixed at their best-fit values from the 5 kb distance. The fitting minimized the squared errors under this constraint.
